# Erythropoietin directly remodels the clonal composition of murine hematopoietic multipotent progenitor cells

**DOI:** 10.1101/2020.04.20.050146

**Authors:** A.S. Eisele, J. Cosgrove, A. Magniez, E. Tubeuf, S. Tenreira Bento, C. Conrad, F. Cayrac, T. Tak, A.M. Lyne, J. Urbanus, L. Perié

## Abstract

The cytokine erythropoietin (EPO) is a potent inducer of erythrocyte development and one of the most prescribed biopharmaceuticals. The action of EPO on erythroid progenitor cells is well established, but its direct action on hematopoietic stem and progenitor cells (HSPCs) is still debated. Here, using cellular barcoding, we traced the differentiation of hundreds of single murine HSPCs, after *ex vivo* EPO-exposure and transplantation, in five different hematopoietic cell lineages, and observed the transient occurrence of high-output Myeloid-Erythroid-megaKaryocyte (MEK)-biased and Myeloid-B-cell-Dendritic cell (MBDC)-biased clones. Single-cell RNA sequencing (ScRNAseq) analysis of *ex vivo* EPO-exposed HSPCs revealed that EPO induced the upregulation of erythroid associated genes in a subset of HSPCs, overlapping with multipotent progenitor (MPP) 1 and MPP2. Transplantation of Barcoded EPO-exposed-MPP2 confirmed their enrichment in Myeloid-Erythroid-biased clones. Collectively, our data show that EPO does act directly on MPP independent of the niche, and modulates fate by remodeling the clonal composition of the MPP pool.

## Introduction

Erythrocytes are the most numerous hematopoietic cells in our body and are constantly renewed^1^. The major inducer of erythroid cell development in steady state and anemic conditions is the cytokine EPO^2^. Recombinant EPO is widely used to treat anemia and is one of the most sold biopharmaceuticals^3^. Previously, EPO was thought to solely target erythroid-committed progenitors and induce their increased proliferation and survival via the EPO receptor (EPOR)^4^. Recently, EPO has also been suggested to act on HSPCs ^5–12^, but the nature of EPÓs effect on HSPC fate remains unresolved despite potential adverse side effects during long-term EPO usage in the clinics and associations of high EPO levels with leukemias^13–15^.

It is well established that EPO can induce HSPCs to cycle, as evidenced by a number of bulk and single cell studies *in vitro* and *in vivo*^5, 8, 9, 11, 12^. It’s role in modulating HSPC fate is less clear however, with a lack of studies that functionally assess HSPC fate at the single cell level *in vivo*, and analyze the direct effect of EPO on HSPCs, and not the effect of the surrounding niche. More specifically, the upregulation of erythroid associated genes in HSPC (LSK CD150^+^ Flt3^-^ CD48^-^) in response to *in vivo* EPO has been observed with bulk transcriptomics^11^, suggesting that HSPCs deviate their fate toward erythroid production. Recent ScRNAseq analysis observed different changes of lineage-associated gene expressions after *in vivo* EPO-exposure of HSPCs^8, 9^. As changes of gene expression do not necessarily result in cell fate modification^14^, functional validations *in vivo* are necessary. In one study such functional validation was performed using bulk transplantation of *in vivo* EPO-exposed HSPCs (LSK CD150^+^ Flt3^-^)^7^. This study showed increased erythroid production and decreased myeloid cell production, concluding that EPO deviates the fate of HSPCs in favor of erythroid production. As EPO has also been shown to target hematopoietic niche cells (osteoblasts and osteocytes^15, 16^ endothelial cells^16, 17^, adipocytes^18, 19^, and mesenchymal stem cells^6, 20^), it remains however still unclear whether EPO acts directly on HSPCs, via their environment, or both.

It is now established that HSPCs encompass cells with different long-term reconstitution capacity after transplantation, as well as a heterogeneous output in terms of quantity (lineage-bias) and type of cells (lineage-restriction)^21–28^. Recently, the HSPC compartment was subdivided into long-term hematopoietic stem cells (LT-HSC) and different multipotent progenitors (MPP) (MPP1-4^29, 30^), with variation around the phenotypic definition^31^. Interestingly, HSPC composition responds to irradiation with HSC transiently self-renewing less and increasing their production of MPP2-3^31^. In the case of EPO, conflicting results suggest that either HSC or MPP respond to EPO ^8, 9, 11^ but the difference in HSPC definition and the lack of functional validation make it difficult to compare these studies. As HSPCs are functionally heterogeneous, and the current phenotypic definition partially capture this heterogeneity, a single cell in vivo lineage tracing approach is needed to assess whether EPO can influence HSPC fate decisions.

To analyze the functional effect of EPO on the differentiation of individual HSPCs (C-Kit^+^ Sca1^+^ CD150^+^ Flt3^-^) removing the effect of the niche, we here utilized cellular barcoding technology which allowed us to trace the progeny of hundreds of single HSPCs *in vivo*. By analyzing cellular barcodes in five mature hematopoietic lineages and HSPCs, we observed transient induction of high-output MEK-biased barcode clones compensated by MBDC-biased clones after ex vivo EPO-exposure. ScRNAseq of *ex vivo* EPO-exposed HSPCs revealed upregulation of erythroid associated genes in a subset of the compartment with overlap to gene signatures of MPP1 (C-Kit^+^ Sca1^+^ Flt3^-^ CD150^+^ CD48^-^ CD34^+^), and MPP2 (C-Kit^+^ Sca1^+^ Flt3^-^ CD150^+^ CD48^+^)^29, 30^ and not of LT-HSCs (C-Kit^+^ Sca1^+^ Flt3^-^ CD150^+^ CD48^-^ CD34^-^). Transplantation of barcoded MPP2 confirmed their enrichment in ME-biased clones in response to EPO. Moreover, the increased contribution of biased HSPC clones to the mature cell lineages after EPO-exposure did not match their frequency in HSPCs, indicating that they were differentiating more than self-renewing, a property associated to multipotent progenitors. The transient effect of EPO on HSPCs further corroborates an action of EPO on MPP1/2 rather than LT-HSCs. Altogether our results are consistent with a model in which perturbations induce clonal remodeling of HSPC contributing to hematopoiesis, with biased MPPs transiently contributing more than LT-HSC. They also demonstrate a direct effect of EPO on MPPs after transplantation with implications for basic HSC research and therapeutic applications in the clinic.

## Results

### EPO-exposure induces biases in single HSPCs

Given the debate surrounding which HSPC subset is responding to EPO, we decided to analyzed the direct effect of EPO on the differentiation of HSPCs defined as C-Kit^+^ Sca1^+^ Flt3^-^ CD150^+^ (encompassing LT-HSC (C-Kit^+^ Sca1^+^ Flt3^-^ CD150^+^ CD48^-^ CD34^-^), MPP1 (C-Kit^+^ Sca1^+^ Flt3^-^ CD150^+^ CD48^-^ CD34^+^), and MPP2 (C-Kit^+^ Sca1^+^ Flt3^-^ CD150^+^ CD48^+^)^29, 30^) at the single cell level by cellular barcoding. To this purpose, we generated a new high-diversity lentiviral barcode library (LG2.2, 18,026 barcodes in reference list), consisting of random 20 nucleotides sequences positioned adjacent to the green fluorescent protein (GFP) gene, enabling the tracking of many individual cells in parallel. Using this LG2.2 library, we labeled single HSPCs (Figure 1-figure supplement 1a) with unique genetic barcodes as previously described^33^, exposed them to EPO (1,000 ng/ml) or PBS for 16 hours *ex vivo* and transplanted around 2,600 cells (Mean 2,684 cells +/- 175 cells) of which around 10% barcoded cells into irradiated mice (Figure 1a). Note that HSPCs kept their sorting phenotype after *ex vivo* culture albeit a slight downregulation of C-Kit^44^ and Flt3 (Figure 1 – figure supplement 1f). At day 30 after transplantation, the earliest timepoint at which HSPCs produce simultaneously erythroid, myeloid and lymphoid cells^45^, barcoded (GFP^+^) erythroblasts (E; Ter119^+^ CD44^+^^34^), myeloid cells (M; Ter119^-^ CD19^-^ CD11c^-^ CD11b^+^), and B-cells (B; Ter119^-^ CD19^+^) (Figure 1 – figure supplement 1b-c, e) were sorted from the spleen and their barcode identity assessed through PCR and deep-sequencing. Note that bone and spleen had similar barcoding profiles (Figure 1 – figure supplement 2). No difference in chimerism was observed between the EPO and control group in the spleen and blood, even when mTdTomato/mGFP donor mice were used to better assess the erythroid lineage (Figure 1b-c). On average, we detected around eighty barcodes per mouse, of which most were detected in several lineages (Figure 1d). Comparison of the numbers of barcodes producing each lineage showed that EPO-exposure resulted in the same number of engrafting and differentiating cells as in control (Figure 1d). Notably, the number of erythroid restricted cells remained stable in the EPO group as compared to control (Figure 1e), indicating that the response to EPO is more complex than a direct instruction of erythroid-restricted HSPCs.

**Figure 1:**
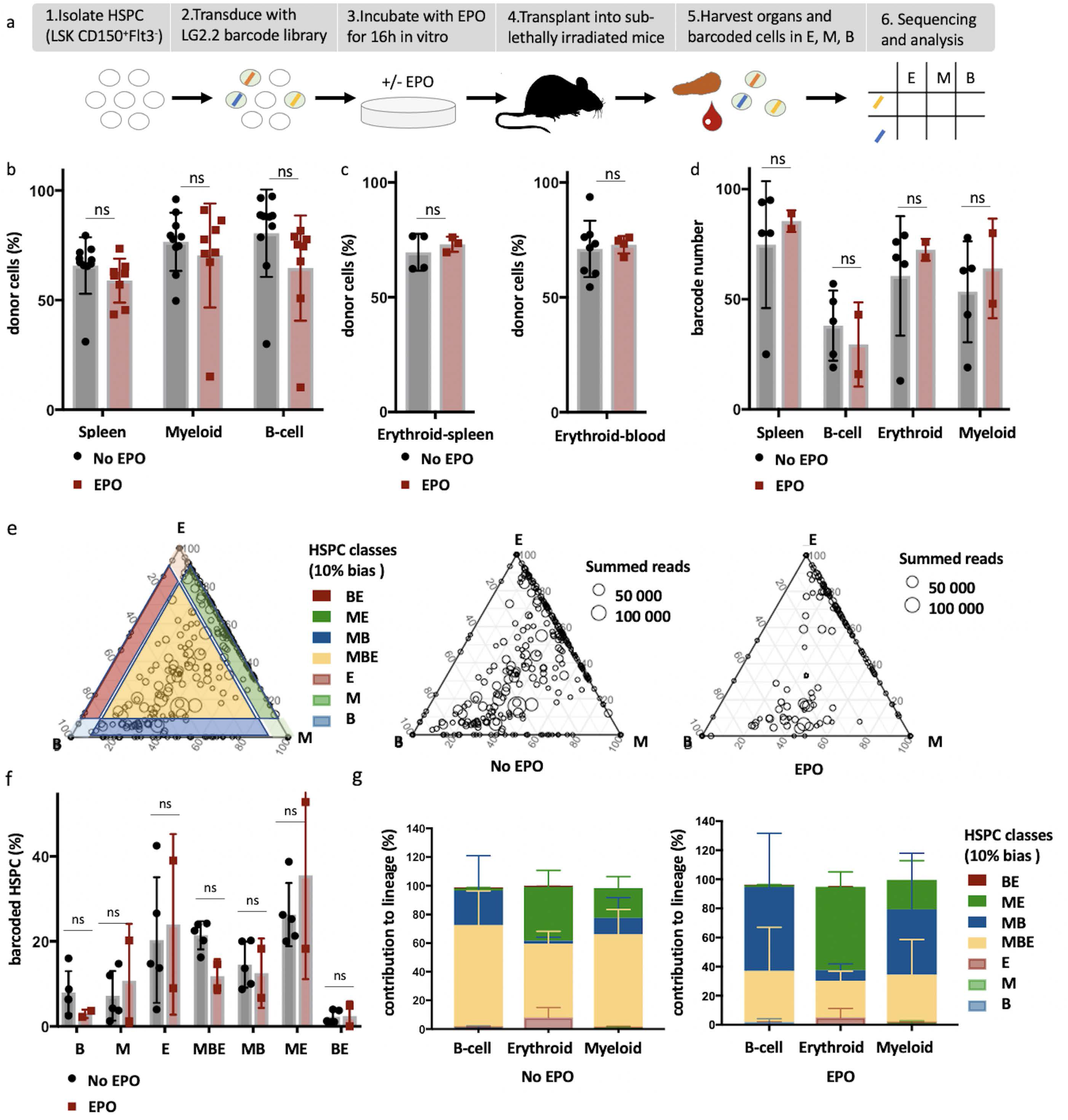
High output ME- and MB-biased clones occur after EPO-exposure and transplantation of HSPCs. **a**, HSPCs were sorted from the bone marrow of donor mice, lentivirally barcoded, cultured *ex vivo* with or without 1,000 ng/ml EPO for 16 h and transplanted into sublethally irradiated mice. At week 4 post-transplantation, the erythroid (E), myeloid (M), and B-cells (B) lineages were sorted from the spleen and processed for barcode analysis. **b**, The percentage of donor derived cells (CD45.1^+^) among the total spleen, myeloid cells (CD11b^+^) or B-cells (CD19^+^) in the spleen of control and EPO group. **c**, To better assess chimerism in erythroid cells mTdTomato/mGFP donor mice were used. The fraction of Tom^+^ cells among erythroid cells (Ter119^+^) in the spleen and blood in control and EPO group. **d**, Number of barcodes retrieved in the indicated lineages at week 4 after transplantation in control and EPO group. **e**, Triangle plots showing the relative abundance of barcodes (circles) in the E, M, and B lineage with respect to the summed output over the three lineages (size of the circles) for control and EPO group. **f**, Percentage of HSPCs classified by the indicated lineage bias, using a 10% threshold for categorization. **g**, Quantitative contribution of the classes as in f to each lineage. Shown are values from several animals (n= 8 EPO, n= 10 control in b, n= 3 EPO, n=4 control in c spleen, n= 4 EPO, n=8 control in c blood collected over 5 different experiments (d-g) n=5 for control and n=2 for EPO group collected over one experiment). For all bar graphs mean and S.D. between mice are depicted. Statistical significance tested using Mann-Withney U test p=0,05 for (b-c). Statistical significance tested by permutation test for different subsets in g (see Table 1). This Figure has four supplements.

**Table 1:**
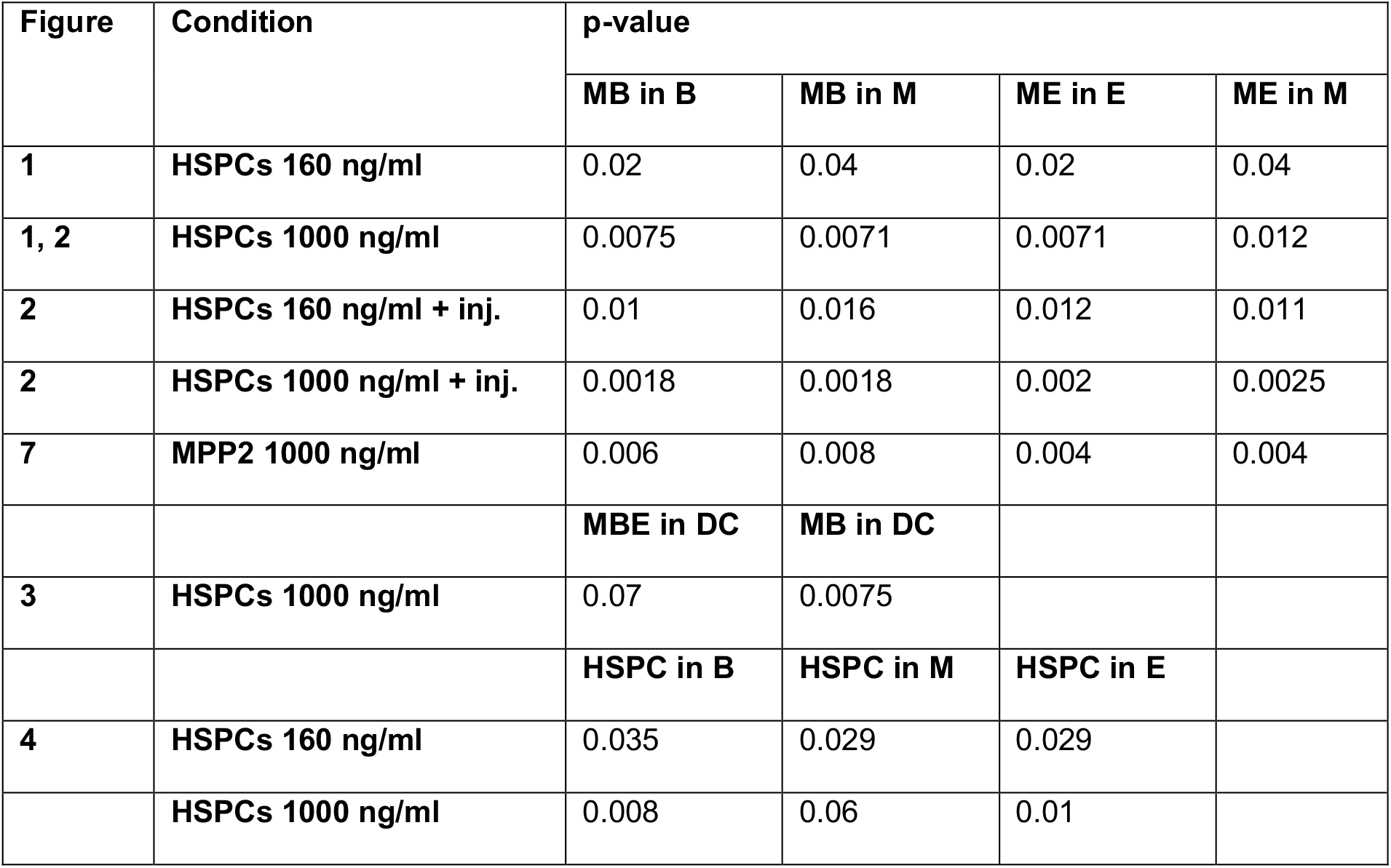
Permutation testing of changes in clonality after transplantation of EPO-exposed HSPCs. Same data as in Figure 1, 2, 3, 4 and Figure 7. HSPCs or MPP2 were cultured with different concentrations of EPO (160 ng/ml or 1,000 ng/ml) for 16h, and when indicated a soluble dose of EPO (133 ug/kg) was injected together with barcoded HSPCs at the moment of transplantation. Barcodes in the erythroid (E), myeloid (M), B-lymphoid (B) lineage, dendritic cell (DC) and HSPCs, were analyzed four weeks after transplantation and categorized by bias using a 10% threshold. For the data of Figure 1, 2 and 7, the output of MB and ME classified barcodes to the B, M, and E lineages was analyzed using a permutation test. For the data of Figure 3, the output of MBE and MB classified barcodes to the DC lineage was analyzed. For the data of Figure 4, the output of barcodes present in HSPCs to the B, M, and E lineages was analyzed using a permutation test. By permutating the mice of control and EPO groups, the random distribution of this output was generated and compared to the real output difference between control and EPO group. A p-value was generated as in^47^

| Figure | Condition | p-value |  |  |  |
| --- | --- | --- | --- | --- | --- |
|  |  | MB in B | MB in M | ME in E | ME in M |
| 1 | HSPCs 160 ng/ml | 0.02 | 0.04 | 0.02 | 0.04 |
| 1, 2 | HSPCs 1000 ng/ml | 0.0075 | 0.0071 | 0.0071 | 0.012 |
| 2 | HSPCs 160 ng/ml + inj. | 0.01 | 0.016 | 0.012 | 0.011 |
| 2 | HSPCs 1000 ng/ml + inj. | 0.0018 | 0.0018 | 0.002 | 0.0025 |
| 7 | MPP2 1000 ng/ml | 0.006 | 0.008 | 0.004 | 0.004 |
|  |  | MBE in DC | MB in DC |  |  |
| 3 | HSPCs 1000 ng/ml | 0.07 | 0.0075 |  |  |
|  |  | HSPC in B | HSPC in M | HSPC in E |  |
| 4 | HSPCs 160 ng/ml | 0.035 | 0.029 | 0.029 |  |
|  | HSPCs 1000 ng/ml | 0.008 | 0.06 | 0.01 |  |

To quantify the effect of EPO on HSPC lineage-biases, barcode-labeled HSPCs were classified based on the balance of their cellular output in the M, B and E lineages. With this classification using a 10% threshold, cells classify for example as ME-biased if they have above 10% of their output in the M and E lineage, and under 10% of their output in the B lineage (Figure 1e other thresholds in Figure 1 – figure supplement 3b). Interestingly, application of this classification revealed that although the proportion of lineage-biased HSPCs in control and EPO group was similar (Figure 1f), their contribution to the different lineages was increased by EPO-exposure (Figure 1g). In the control group, balanced HSPCs (MBE) produced the majority of all lineages, as previously published^46^. In the EPO group, ME- and MB-biased clones produced most cells of the analyzed lineages (Figure 1g). ME-biased HSPCs produced the majority of erythroid cells (57% +/- 10%), MB-biased HSPCs produced the majority of B-cells (58% +/- 36%), and ME- and MB-biased clones contributed the majority of myeloid cells (MB-biased 45% +/- 38% and ME-biased 20% +/- 13%, together 65% +/- 25%). To test the significance of this effect, we used a permutation test, which compares the effect size between control and EPO group to the one of all random groupings of mice^47^. The contributions of the ME- and of the MB-biased HSPC classes to the different lineages were significantly different in EPO and control groups (Table 1). These results were reproduced in an additional experiment (Figure 1 – figure supplement 3c-f, Supplementary File 1). A lower EPO concentration (160 ng/ml) as well as an additional single injection of EPO (133 ug/kg) during transplantation gave similar results (Figure 2, Figure 2 – figure supplement 1a, Table 1). Also at six weeks post transplantation, similar results were obtained (Figure 1 – figure supplement 4, Supplementary File 1). In summary, *ex vivo* EPO priming of HSPCs modified the output balance of HSPCs rather than the number of lineage-restricted and -biased cells. Balanced clones produced a smaller percentage of the mature cells; ME-biased HSPCs produced most of the erythroid cells and MB-biased HSPCs produced most of the B cells.

**Figure 2:**
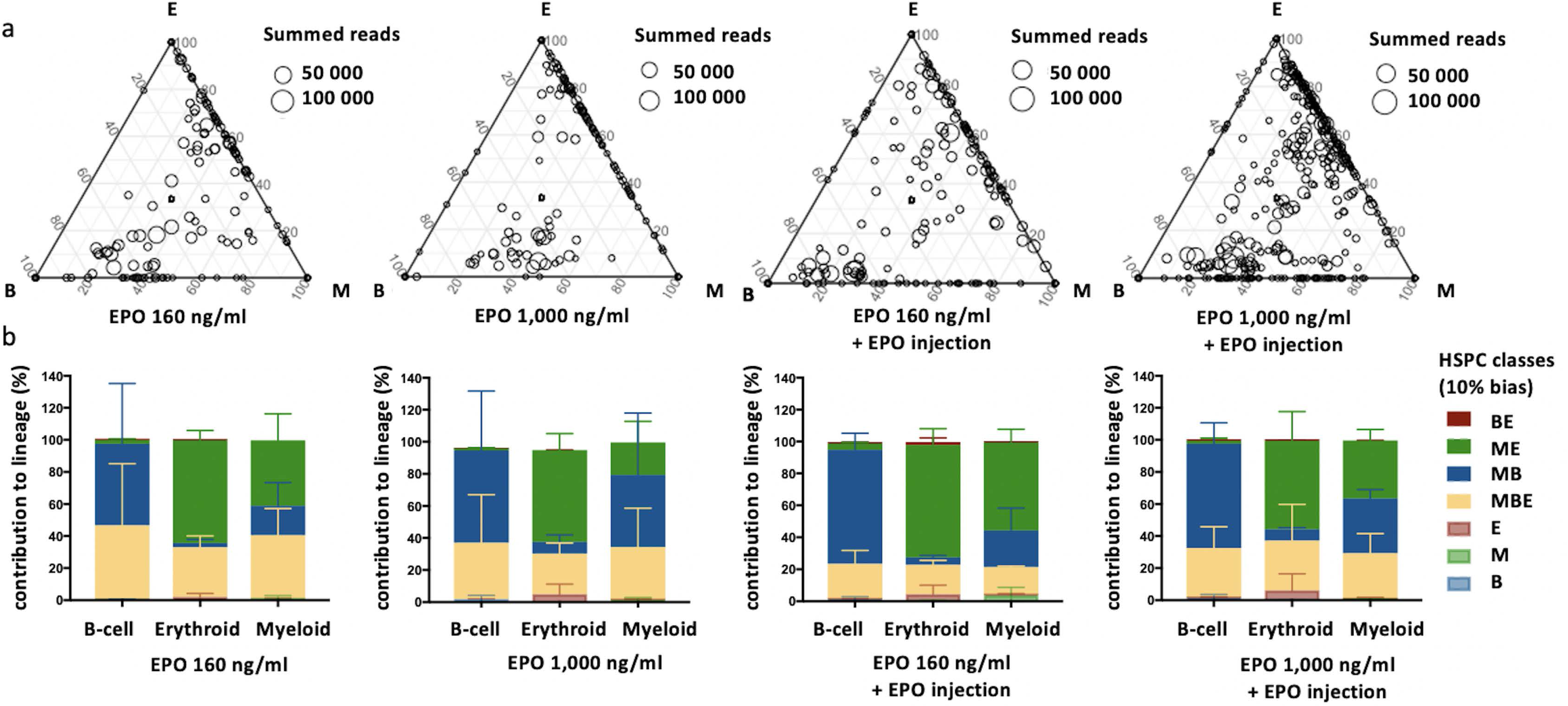
Effect of different EPO concentrations on HSPC clonality after transplantation. Same protocol as in Figure 1 but HSPCs were cultured with different concentrations of EPO (160 ng/ml or 1,000 ng/ml) for 16 h, and when indicated a single dose of EPO (133 ug/kg) was injected together with barcoded cells at the moment of transplantation. **a**, Triangle plots showing the relative abundance of barcodes (circles) in the erythroid (E), myeloid (M), and B-lymphoid (B) lineage with respect to the summed output over the three lineages (size of the circles) for the different experimental groups as indicated. **b**, The percentage of each lineage produced by the barcodes categorized by bias using a 10% threshold. Shown are values from several animals (n=2 for 160 ng/ml, 1,000 ng/ml, and 160 ng/ml + EPO injection, n=4 for 1,000 ng/ml + EPO injection (collected over 4 different experiments)). For all bar graphs mean and S.D. between mice are depicted. Statistical significance tested by permutation test for different subsets in b (see Table 1). This figure has one supplement.

### Contribution of ME- and MB-biased HSPCs to the DC and MkP lineage

To further characterize the cells produced by the ME-biased and MB-biased HSPCs, we repeated our experimental setup including the analysis of the megakaryocyte and dendritic cell (DC) lineages (Figure 3). Megakaryocyte Progenitors (MkP) were chosen as proxy for the production of platelets which are not suitable for barcode analysis. Barcoded (GFP^+^) DCs (DC; Donor Ter119^-^ CD19^-^ CD11c^+^ CD11b^-^) and MkP (MkP; C-Kit^+^ Sca-1^-^ CD150^+^ CD41^+^) (Figure 1 – figure supplement 1c-e) were sorted together with M, E, and B cells, 4 weeks after transplantation of control or EPO-exposed HSPCs (1,000 ng/ml). In both groups, the majority of clones produced also DCs (Figure 3a). In the control group, balanced HSPCs produced the majority of DCs (65% +/- 9%) (Figure 3b). However, in the EPO group, balanced HSPCs decreased their contribution to the DC lineage (36% +/-25%) and MB-biased HSPCs significantly increased their contribution (86% +/- 43% EPO vs 22% +/- 11% control group) (Figure 3b, Table 1), thus, they were MBDC-biased HSPCs. In contrast the ME-biased HSPCs produced few DCs in both groups (Figure 3b), indicating that ME-biased HSPCs are restricted both in their B and DC production compared to the M and E production.

**Figure 3:**
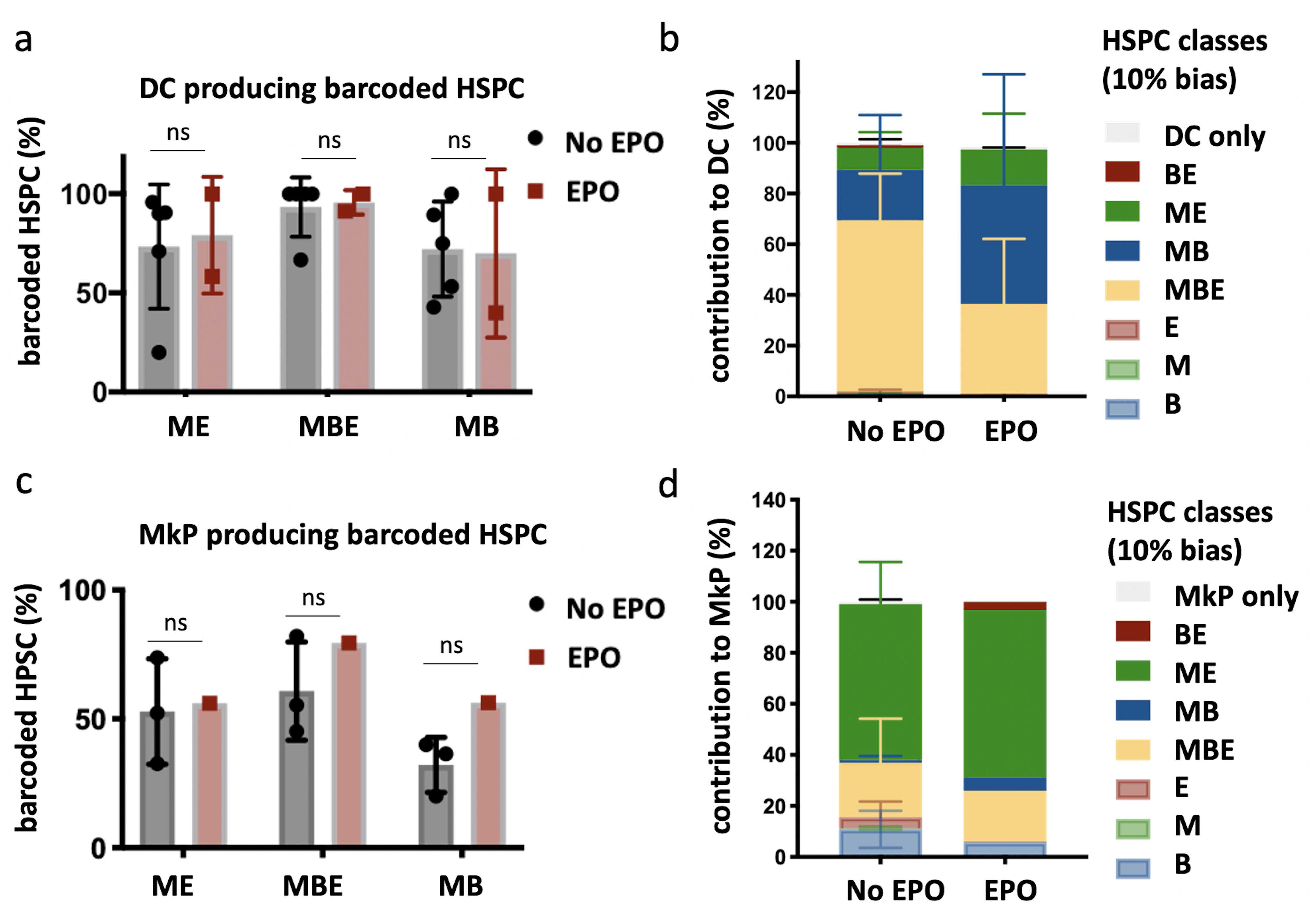
Production of Dendritic Cells (DC) and Megakaryocyte Progenitors (MkP) by HSPCs after EPO-exposure and transplantation. In addition to the analysis of barcodes in the erythroid (E), the myeloid (M), and the B-cell (B) lineage, the DC lineage in spleen and MkP in bone marrow were added. **a**, Percentage of barcoded HSPCs producing DC in the different HSPC categories (classification as in Figure 2 based on the M, E, and B lineage only using a 10% threshold. The DC only category was added). **b**, The percentage of the DC lineage produced by the barcodes categorized by bias as in a. **c-d**, Representations as in a-b for barcode detection in MkP. Data is derived from a cohort with detailed myeloid sorting. The myeloid lineage was merged according to the percentage of total donor myeloid each subset contributed as in Figure 2 – supplement 1a to allow classification as in a-b based on the M, E, and B lineage only using a 10% threshold. The MkP only category was added. Shown are values from several animals (a-b, n=5 for control and n=2 for EPO group, c-d, n=3 for control and n=1 for EPO group (collected over two experiments)). For all bar graphs mean and S.D. between mice are depicted. Statistical significance tested using Mann-Withney U test p=0,05 for (a,c). Statistical significance tested by permutation test for different subsets in b (see Table 1). This figure has one supplement.

The majority of the MkP production came from the ME-biased HSPCs in both groups (58% +/- 21% control and 55% +/- 14% EPO group, Figure 3c-d), indicating that ME-biased HSPCs were also MkP-biased HSPCs (thus MEK-biased). We did not detect a high contribution of MkP-restricted HSPCs^24, 25, 48^ to the MkP lineage (Figure 3d). Finally, as high EPO-exposure has been linked to changes in macrophage numbers^49–, 56^ we analyzed the contribution of control and EPO-exposed HSPCs to the myeloid lineage in more detail, but could not detect changes in the percentage of the different myeloid subsets produced (Figure 3 – figure supplement 1a-c).

### Effect of EPO on short-term HSPC self-renewal

In light of previous studies which suggested changes in HSPC proliferation after *in vivo* EPO-exposure^5, 8, 9, 11, 12^, we next explored if the short-term self-renewal capacity of HSPCs was impacted. To this end, we analyzed barcodes in bone marrow HSPCs in addition to the spleen E, M, and B lineages at week 4 after transplantation of control or EPO-exposed HSPCs (160 and 1,000 ng/ml) (Figure 4). We reasoned that barcodes of HSPCs differentiating and short-term self-renewing (dividing to give rise to other HSPCs) after transplantation are detected in both compartments, while detection in only HSPCs or mature lineages indicates a prevalence of short-term self-renewal or differentiation respectively. Most of the barcodes detected in HSPCs overlapped with barcodes in the mature cells (Figure 4b left) in both the control and two EPO groups, showing that most of the transplanted cells had given rise to other HSPCs and differentiated irrespective of the treatment. Some barcodes were only detected in mature cells (Figure 4b right), indicating that some HSPCs had only differentiated or were below the limit of detection. These HSPCs were equally abundant in the control and two EPO groups (Figure 4b right).

**Figure 4:**
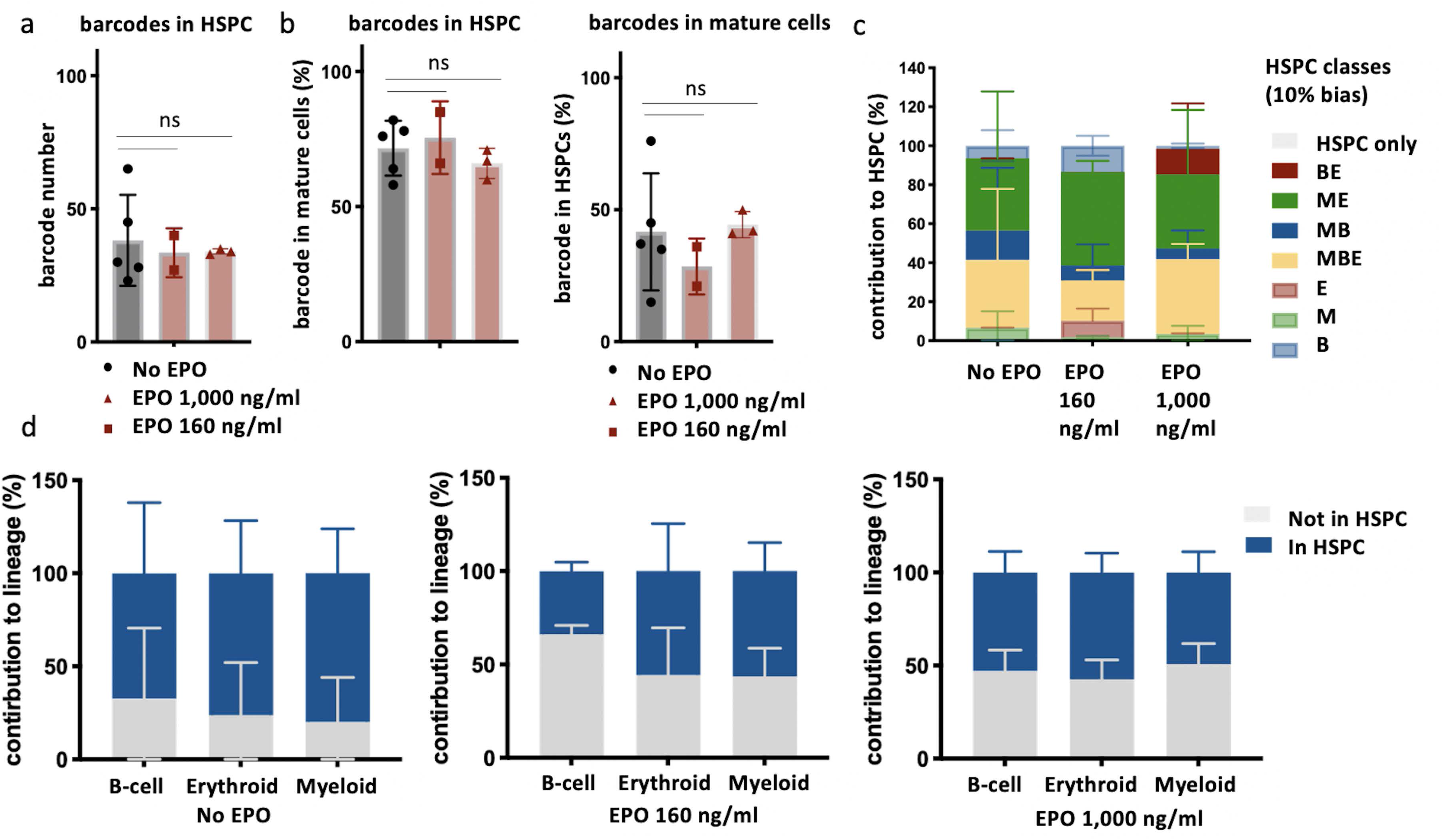
Overlap of barcodes in HSPCs and mature cells after transplantation of EPO-exposed HSPCs. Same protocol as in Figure 1 but HSPCs were cultured with two different concentrations of EPO (160 ng/ml or 1000 ng/ml) for 16 h. In addition, HSPCs were sorted and subjected to barcode analysis. **a**, The total number of barcodes found back in HSPCs. **b**, The percentage of barcodes in the mature cell subsets also detected in HSPCs and the percentage of barcodes in HSPCs also detected in mature cells. **c**, The percentage of the HSPC lineage contributed by barcodes categorized by bias as in Figure 2 based on the M, E, and B lineage using a 10% threshold. **d**, The percentage of each lineage produced by the barcodes color coded for presence (blue) and absence (grey) in HSPCs. Shown are values from several animals (n=5 for control, n=2 for EPO 160 ng/ml group and n=3 for EPO 1,000 ng/ml group). For all bar graphs mean and S.D. between mice are depicted. Statistical significance tested using Mann-Withney U test p=0,05 for (a-b). Statistical significance tested by permutation test for different subsets in d (see Table 1).

To analyze if different lineage biases correlated to different short-term self-renewal capacity, we analyzed the proportion of biased HSPC classes, as previously defined, within the HSPC compartment (Figure 4c). In the control group, balanced and ME-biased HSPCs contributed most to the HSPC reads (34% +/- 36% MBE and 37% +/- 34% ME-biased HSPCs), while barcodes of MB-biased HSPCs contributed less (15% +/- 32%) (Figure 4c), a trend that has been previously described^25, 57, 58^. Surprisingly, the pattern of contributions of different biased HSPC subsets to HSPC reads was unchanged in the EPO groups (Figure 4c), implying that the extent of short-term self-renewal was unchanged after *ex vivo* EPO-exposure.

To study if the increased production of cells by the ME- and MB-biased HSPCs to the mature cells observed after *ex vivo* EPO-exposure (Figure 1-3) correlated with short-term self-renewal capacity of HSPCs, we analyzed the contribution of barcodes detected or not in HSPCs to the E, M, and B lineages (Figure 4d). In the control group, the majority of mature cells were derived from barcodes also present in HSPCs. However, in both EPO groups, the contribution of barcodes detected in HSPCs to mature cells was significantly lower (Figure 4d, Table 1) implying that the increased contribution of biased HSPC classes to the mature cell lineages after *ex vivo* EPO-exposure was most likely caused by cells differentiating more than short-term self-renewing.

### EPO-exposure induces an erythroid program in a subgroup of HSPCs

To further characterize the effect of EPO-exposure on HSPCs, we performed scRNAseq of barcoded C-Kit^+^ Sca1^+^ Flt3^-^ CD150^+^ cells after *ex vivo* culture in medium supplemented with EPO or PBS using the 10X Genomics Chromium platform. 1,706 cells from control and 1,595 cells from the EPO group passed our quality control. To compare the HSPCs injected with non-cultured hematopoietic cells, we generated a reference map of 44,802 C-kit^+^ cells from^42^ and used published signatures as detailed in the supplementary materials and methods^31, 43^ to annotate this map (Figure 5 – figure supplement 1a,b and e,f). Projection of our single cell data on this map showed that both the control and the EPO-exposed HSPCs similarly overlapped with non-MPP4 LSK cells, according to their sorting phenotype (Figure 5 – figure supplement 1e,f). These results indicate that neither the *ex vivo* culture itself nor the EPO treatment dramatically affected the global identity of the sorted HSPCs.

When comparing the EPO and control group, we found 1,176 differentially expressed genes (Figure 5a and Supplementary File 2) and this number was significantly higher than the number expected due to chance (p-value=0,01) as assessed by permutation testing. Among the most upregulated genes in the EPO-exposed HSPCs, were genes with erythroid association as *Hbb-bs*, *Erdr1*, *Wtap*, *Kmt2d*, or *Nfia*^59^, and GATA1 targets (*Abhd2*, *Cbx3*, *Kdelr2*, *Pfas*), cell cycle related genes (*Tubb5*, *Hist1h2ap*) as well as genes previously described to be induced in HSPCs after *in vivo* EPO-exposure, such as *Bmp2k*^6^ and *Ifitm1*^8^ (Figure 5a). Genes involved in stem cell maintenance, such as *Serpina3g*, *Mecom, Txnip*, *Meis1*, *Pdzk1ip1*^8^, *Sqstm1*^60^, *Smad7*^61^, *Aes*^62^, were among the most downregulated genes in the EPO-exposed HSPCs (Figure 5a).

**Figure 5:**
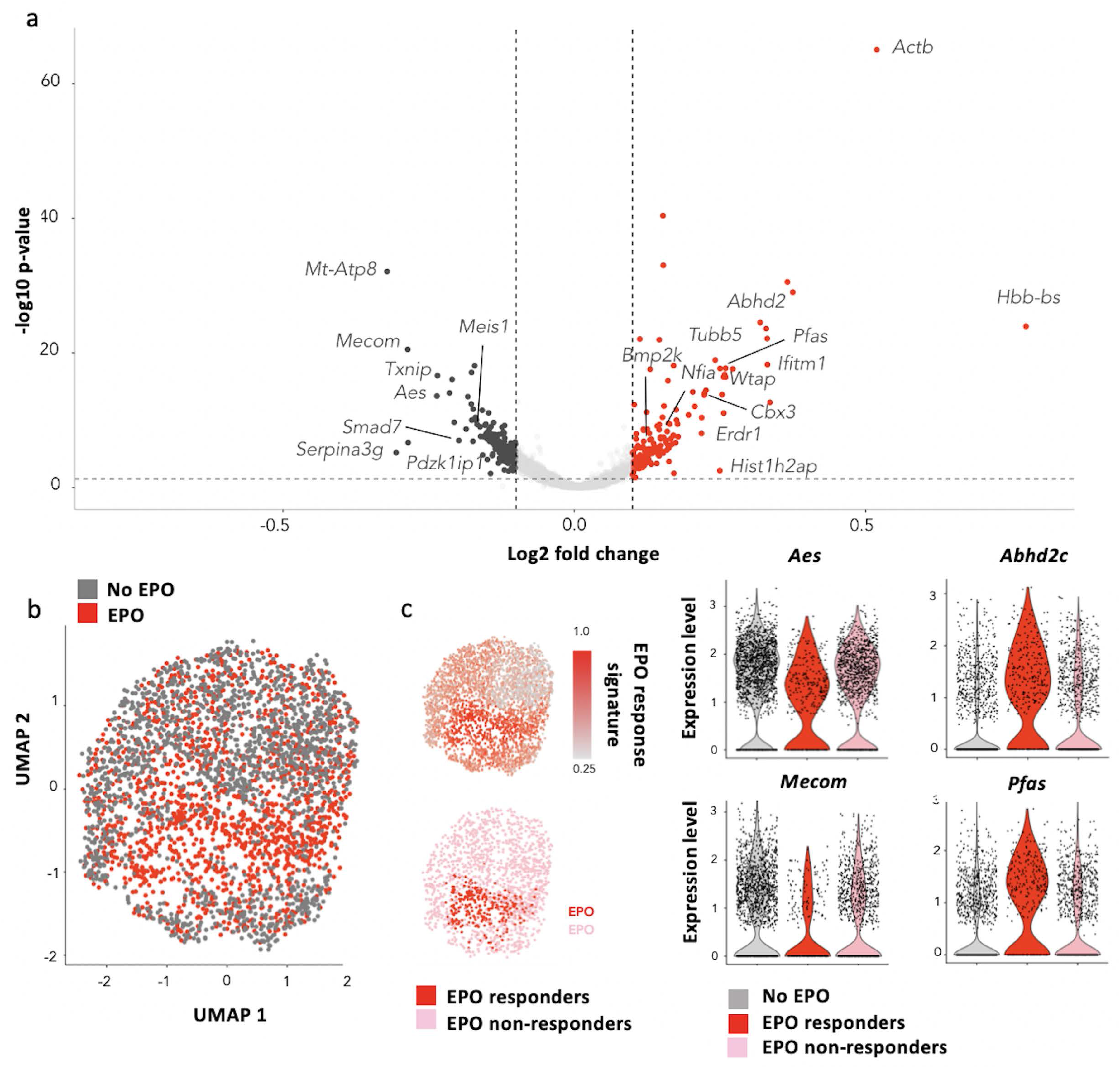
Characterization of EPO-exposed HSPCs by scRNAseq. HSPCs were sorted, barcoded and cultured *ex vivo* with or without 1,000 ng/ml EPO for 16 h, and analysed by scRNAseq using the 10X Genomics platform. 1,706 cells from control and 1,595 cells from EPO group passed quality control **a**, Volcano plot of log2 fold change of the differentially expressed genes between control and EPO-exposed cells versus the adjusted p-value. Genes of interest are annotated. Differentially expressed genes were used to defined an EPO response signature. **b**, UMAP visualization of the EPO-exposed and control HSPCs. **c**, The level of expression in the EPO-exposed HSPCs of the genes in the EPO response signature (top), and definition of the EPO-responder and non-responder subgroups using the 90th percentile expression of the EPO response signature from c (bottom). **d**, The expression of the indicated genes in the control, EPO-responder and non-responder subgroups as defined in c. Genes that are significantly upregulated in the EPO-responder group, when compared to control and non-responder groups. Differential expression was assessed using a logistic regression testing approach, as implemented in Seurat. This figure has two supplements and correspond to one 10X experiment of a pool of 8 mice.

As our cellular barcoding data suggests that single HSPCs differ in their response to EPO, we assessed the heterogeneity of EPO responses at the transcriptomic level. UMAP-based visualization of the data suggested that a subgroup of EPO-exposed cells was transcriptomically distinct (Figure 5b), independently of the number of PCA components and genes used in the analysis (Figure 5 – figure supplement 1c). To test this observation, we defined an EPO response signature based on differentially expressed genes between the EPO and control group. Plotting the expression of the EPO response signature at the single cell level showed that the majority of the transcriptomic differences between the control and EPO group were indeed driven by this small subgroup of cells (Figure 5c). Reasoning that this subgroup contains the cells directly responding to EPO, we defined as EPO-responders, cells in the 90^th^ percentile of EPO response signature expression (Figure 5c) for subsequent analysis. Importantly, unsupervised clustering analysis of the data (Figure 5 – figure supplement 2a-b) showed similar results. The genes encoding EPOR, as well as the alternative EPO receptors EphB4, CD131, CRFL3 were equally expressed between the EPO-responders, non-responders and control groups (Figure 5 – figure supplement 1d). Reasoning that the EPO-responders correspond to MEK-biased HSPCs, we also looked for potential MBDC-biased HSPCs but could not detect a subgroup of cells with upregulation of lymphoid associated genes, suggesting that the MBDC-bias is not a direct effect of EPO-exposure but more an indirect effect. In summary, the scRNAseq analysis corroborated our functional barcoding data, showing that a subset of HSPCs can respond directly to EPO stimulation.

### EPO-responder HSPCs overlap with MPP1 and MPP2 signatures

As our barcode analysis suggested that the effect of direct EPO-exposure on HSPCs is caused by cells differentiating more than self-renewing, we next wanted to assess which of the HSPC subsets are the EPO responders in our scRNAseq dataset. We annotated the UMAP-based visualization of our data with published signatures of the HSC (dormant HSCs^40^ and LT-HSC^41^), MPP1^40^ and MPP2^31^ subsets included in our HSPC gate, and analyzed its overlap with the previously defined EPO-responder and non-responder cells (Figure 6a,b). Relative to the control group and non-responders of the EPO-group, the EPO-responders had a reduced expression of HSC gene signatures and increased expression of MPP1 and MPP2 signatures (Figure 6a,b). An annotation of the reference map generated from data of^42^ likewise showed a low overlap of EPO-responders with the most quiescent HSC subsets (Figure 6c,d). The independent analysis using unsupervised clustering further supported this result (figure 5 – figure supplement 2c,d). All in all, our scRNAseq analysis implied that, in line with our barcoding results, the HSPCs directly reacting to EPO are most likely multipotent progenitor cells of the MPP1 and MPP2 subsets.

**Figure 6:**
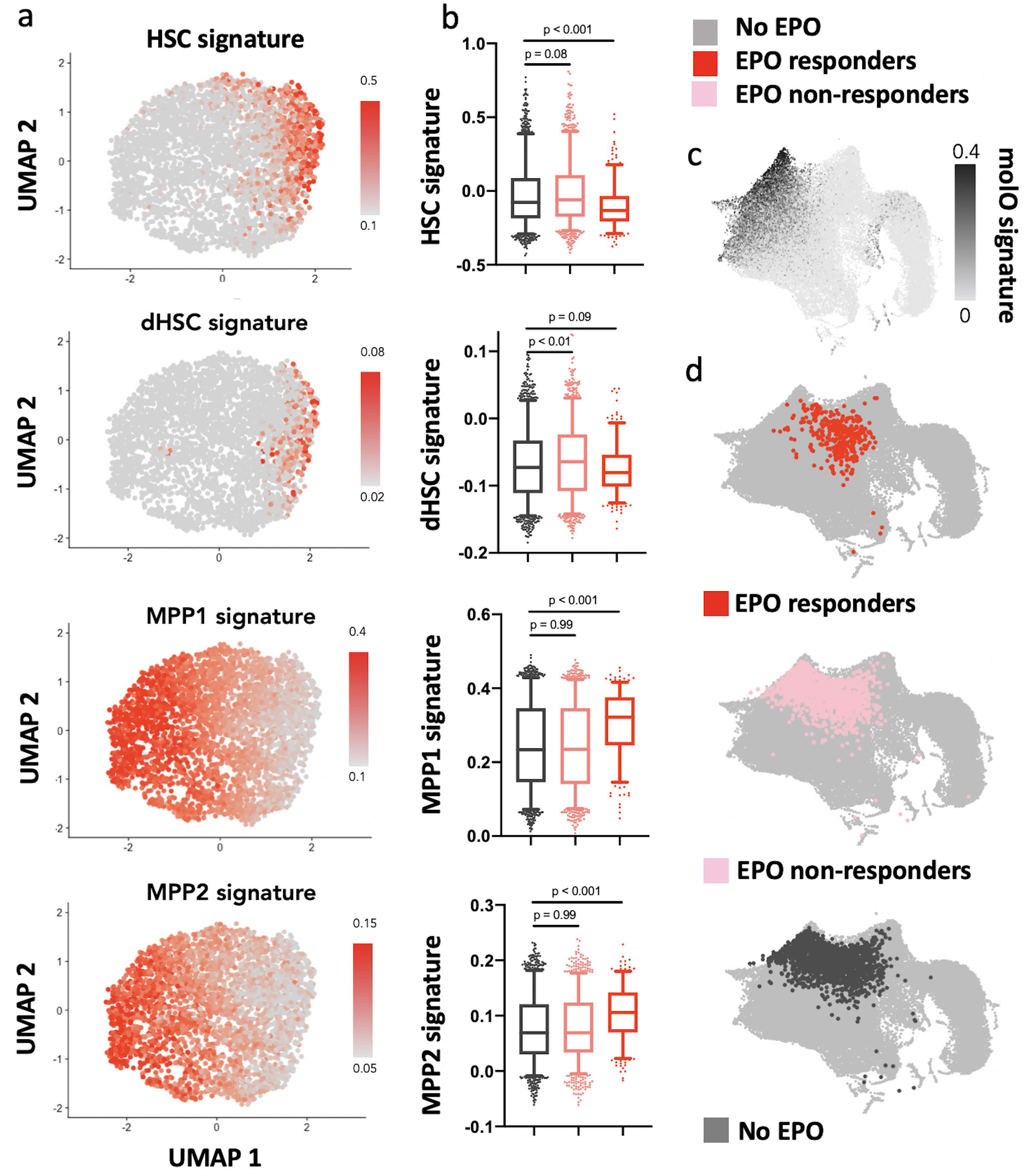
EPO-responders are multipotent progenitors, not HSCs. Same protocol as in Figure 5**. a,** Expression of published gene-signatures of HSCs (dormant HSC^40^, molecular overlap (molO) HSC signature^41^) and MPPs (MPP1^40^-2^31^) across the entire dataset (see Methods) **b**, Expression of the signatures from 6a, across control, non-responder, and EPO-responder groups as defined in Figure 5c. Statistical comparisons made using a Kruskal Wallis Test with a Dunns multiple comparisons post-hoc test **c**, Expression of the molO HSC signature on the published reference map^42^. **d**, Nearest-neighbor mapping of control, EPO-responder and non-responder cells onto the published reference map^42^.

### EPO-exposure induces ME-biases in single MPP2

To confirm that MPP2 are a subset within HSPCs reacting directly to EPO as predicted by the scRNAseq analysis, we transplanted barcoded control or EPO-exposed (1,000 ng/ml) MPP2 together with unbarcoded CD48^-^ HSPCS (C-Kit^+^ Sca1^+^ Flt3^-^ CD150^+^ CD48^-^), (Figure 7 – figure supplement 1a) and analyzed their barcoded progeny in the E, M, and B lineages of the spleen at week 4 after transplantation (Figure 7). We found an equivalent engraftment as for the entire HSPC compartment and no difference between the EPO-treated and the control group (Figure 7a-b). Applying the same classification as in Figure 1 to quantify the effect of EPO on MPP2 lineage-biases, we observed that, as for the whole HSPC compartment, ME-biased cells contributed more to the M and E lineages (Figure 7d-e, other threshold in Figure 7 – figure supplement 1c). Similarly to our data on whole HSPC compartment (Figure 1g) the proportion of the differently biased-MPP2 was similar between control and EPO group (Figure 7c). This data confirms that the MPP2 population is enriched in HSPCs responding to EPO.

**Figure 7:**
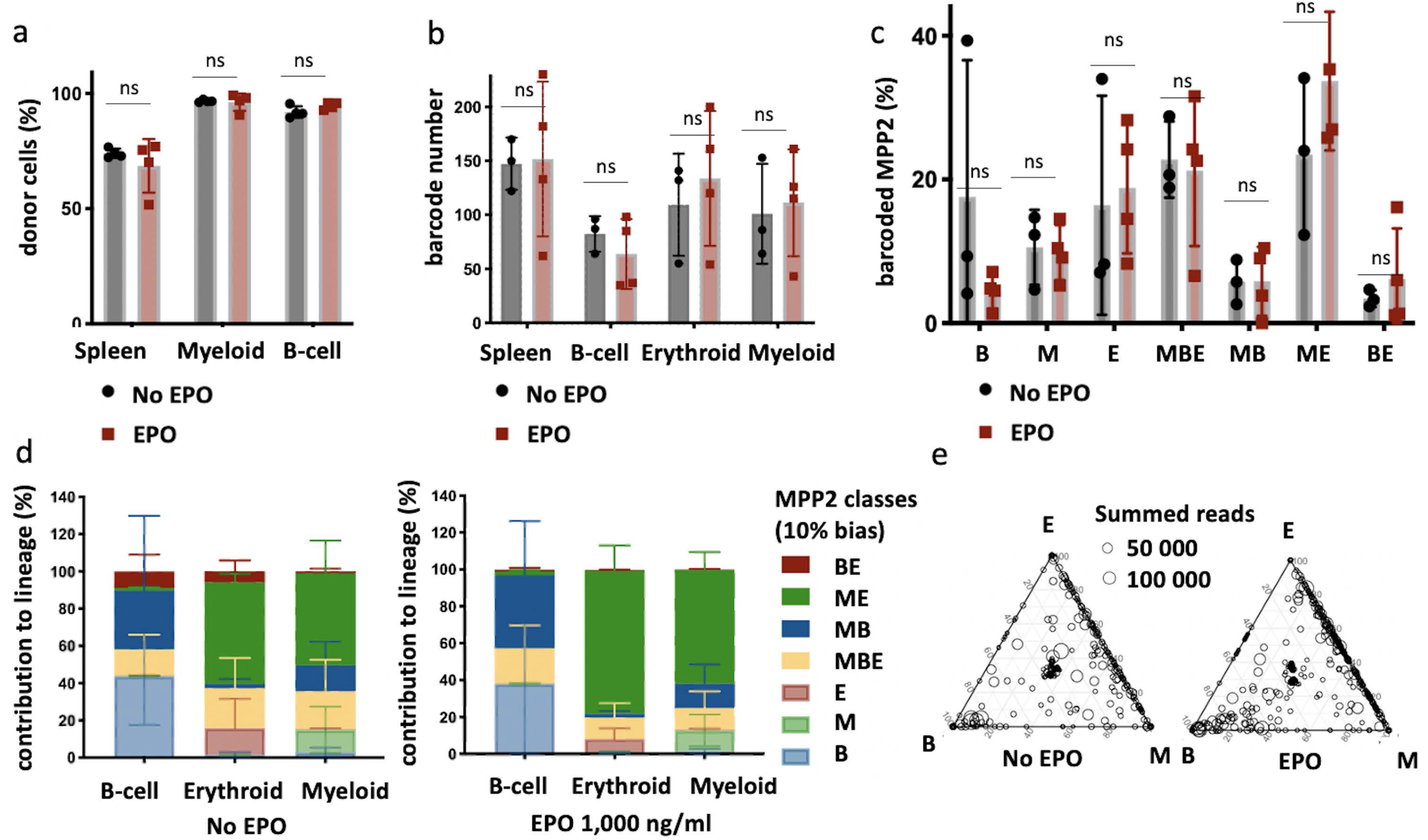
MPP2 are enriched for ME-biased clones after EPO-exposure and transplantation. MPP2 and CD48^-^ HSPCs were sorted from the bone marrow of donor mice, MPP2 were lentivirally barcoded, and both populations cultured *ex vivo* with or without 1,000 ng/ml EPO for 16 h. After the culture, barcoded MPP2 and un-barcoded CD48^-^ HSPCs were mixed and transplanted into sublethally irradiated mice. At week 4 post-transplantation, the erythroid (E), myeloid (M), and B-cells (B) lineages were sorted from the spleen and processed for barcode analysis. **a**, The fraction of donor cells among the indicated cell types in spleen. **b**, Barcode number retrieved in the indicated lineage at 4 weeks after transplantation in control and EPO 1,000 ng/ml group. **c**, Percentage of MPP2s classified using a threshold of 10% in experimental groups as indicated. **d**, The percentage of each lineage produced by the MPP2 barcodes categorized by bias using a 10% threshold. **e**, Triangle plots showing the relative abundance of barcodes (circles) in the erythroid (E), myeloid (M), and B-lymphoid (B) lineage with respect to the summed output over the three lineages (size of the circles). Shown are data from several mice (n=3 for control and n=4 for EPO group). For all bar graphs mean and S.D. between mice are depicted. Statistical significance tested using Mann-Withney U test p=0,05 for (c-e). Statistical significance tested by permutation test for different subsets in a (see Table 1). This figure has one supplement.

### Transient effect of EPO-exposure

Finally, we reasoned that if EPO directly acts on multipotent progenitors MPP1/2 with short reconstitution capacity after transplantation rather than long-term repopulating HSC, then the EPO effect should be transient. To test this hypothesis, we repeated the experiment and analyzed barcodes in the E, M, and B lineages at 4 months after transplantation of control or EPO-exposed HSPCs (160 and 1,000 ng/ml) (Fig. 8). In the control group, as reported before^63^, the chimerism at 4 months was higher and the number of barcodes detected was lower than at one month post-transplantation (Fig. 8c-d and 1b-d). We detected no significant changes in the clonal output of HSPCs between control and EPO group at this timepoint (Fig. 8e), with the majority of cells in all lineages produced by balanced HSPCs (Fig. 8a-b), implying that the effect of direct EPO-exposure on HSPCs is transient. This confirms that the effect of direct EPO-exposure on HSPCs is likely caused by multipotent progenitor cells with a short reconstitution capacity after transplantation.

**Figure 8:**
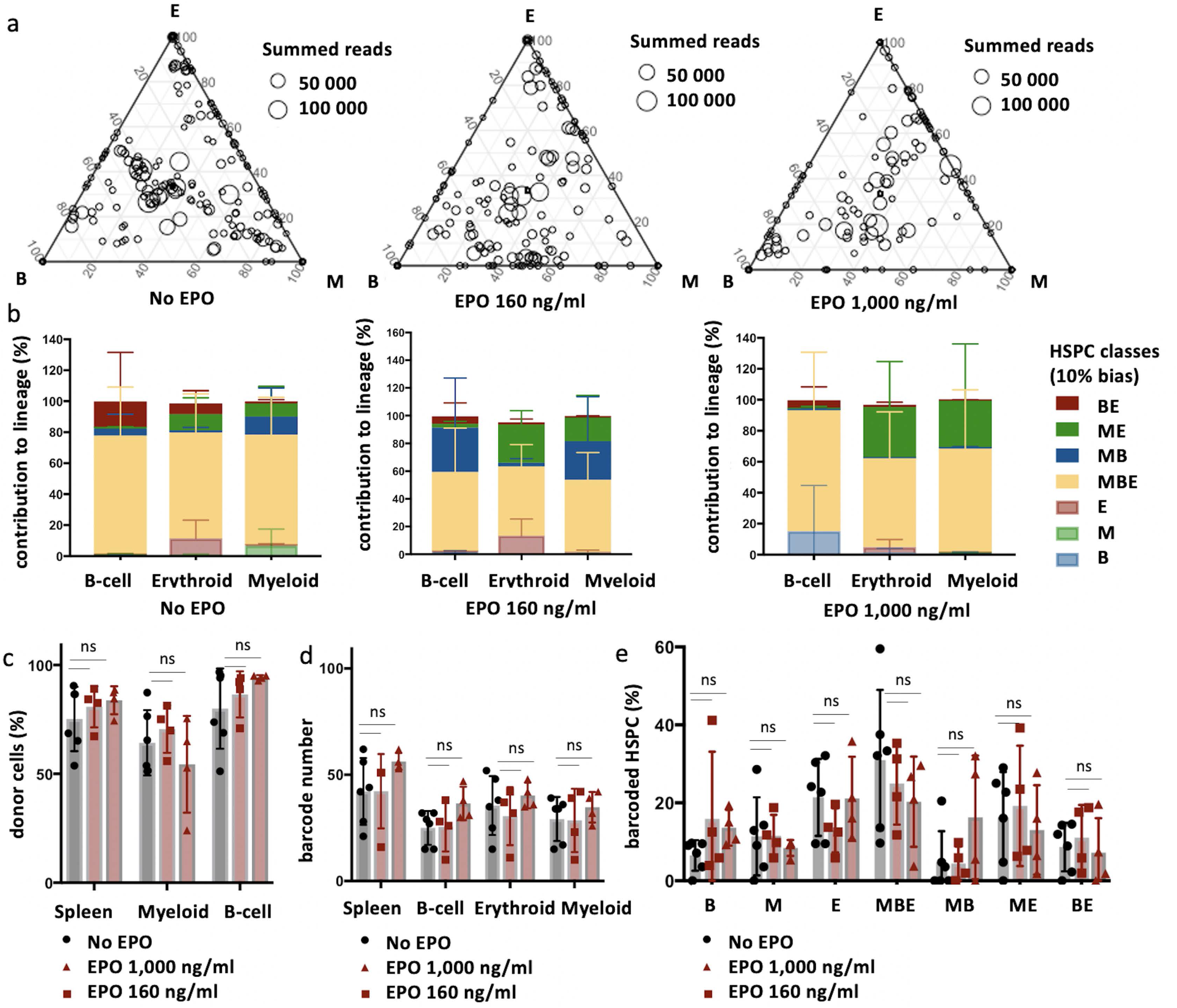
The effect of EPO on HSPC clonality after transplantation is transient. Same protocol as in Figure 1, but barcodes in the E, M, and B lineage in spleen of individual mice sacrificed at month 4 post-transplantation were analyzed. **a**, Triangle plots showing the relative abundance of barcodes (circles) in the erythroid (E), myeloid (M), and B-cell (B) lineage with respect to the summed output over the three lineages (size of the circles) for the different experimental groups as indicated. **b**, The percentage of each lineage produced by the barcodes categorized by bias using a 10% threshold. **c**, The fraction of donor cells among the indicated cell types in spleen. **d**, Barcode number retrieved in the indicated lineage at month 4 after transplantation in control, EPO 160 ng/ml, and EPO 1,000 ng/ml group. **e**, Percentage of HSPCs classified using a threshold of 10% in experimental groups as indicated. Shown are data from several mice. (c, n=5 for control and n=4 for each EPO group, (a-b, d-e) n=6 for control and n=4 for each EPO group (collected over two experiments)). For all bar graphs mean and S.D. between mice are depicted. Statistical significance tested using Mann-Withney U test p=0,05 for (c-e).

## Discussion

EPO is a key regulator of hematopoiesis, and is classically considered to support the proliferation and survival of erythroid committed progenitors. By analyzing the *in vivo* fate of hundreds of EPO-stimulated vs untreated transplanted (c-Kit^+^ Sca1^+^ Flt3^-^ CD150^+^) HSPCs at the single-cell level, we established that EPO can change HSPC differentiation in the absence of an EPO-stimulated bone marrow microenvironment. Collectively our results yield 2 important conclusions: (i) EPO has a direct effect on HSPCs, that is not solely due to the effects of EPO on the surrounding niche and (ii) EPO directly remodels the clonal composition of HSPC by inducing fate biased MPP and reducing the output of HSC.

Specifically, we observe that EPO induced MEK-biased (ME) and MBDC-biased (MB) HSPCs that produced the majority (>60%) of mature hematopoietic cells at four and six weeks after transplantation. In contrast, balanced HSPCs (MBE) had a reduced output of mature cells in response to EPO. The increased erythroid-associated gene signature in a subset of HSPCs after *ex vivo* EPO-exposure suggests that EPO directly induces high output MEK-biased HSPCs, which is indirectly compensated for by the occurrence of high-output of MBDC-biased HSPCs to maintain a balanced production of hematopoietic cells.

These biased clones had a higher propensity to differentiate than to self-renew, and their response to EPO was transient, suggesting that EPO-responsive cells are multi-potent progenitors, and not LT-HSCs. This is supported by transcriptomic analysis showing that EPO-responders express the MPP1/MPP2 gene-signatures. Transplantation of barcoded EPO-exposed MPP2 confirmed their enrichment in ME-biased clones in response to EPO.

Similar to studies that assessed the effect of high systemic EPO-exposure ^8,9,10,11^, we found that multipotent progenitors, not HSC, are responding to EPO. The occurrence of myeloid, megakaryocytic and erythroid gene expression in MPP1 after bleeding ^9^ is in line with our findings. Previously, long-term EPO exposure in Tg6 transgenic mice did however not change the in vitro differentiation outcome of MPP2 ^11^. Furthermore, we did not detect a fate deviation toward erythroid production at the expense of myeloid production as seen for in vivo EPO-exposed HSPCs after transplantation^7^. These differences could be due both to the duration and route of EPO exposure as well as the indirect effects of systemic EPO-exposure through other cells, for example from the bone marrow niche.

In addition, our data shows that direct cytokine-stimulation leads to a clonal remodeling of the HSPC compartment, with a transient increase in the contribution of fate-biased MPPs. Without longitudinal barcoding data within the same animal, we cannot distinguish if EPO is transiently changing the fate and outcome of the same HSPCs over time or if EPO is pushing the differentiation of some HSPCs that will be replaced by more balanced and stable HSPCs. Different studies have suggested that the behavior of transplanted HSPC differ from native HSPC^64, 65^. Transplantation seems to favor the long-term output from HSC whereas steady state hematopoiesis is maintained more by MPPs contribute than LT-HSC^64,65,66^. Our work is in line with a model in which MPP are a highly malleable cell population that can rapidly respond to changing demands for new cells, such as transplantation or infection^31^.

The direct effect of EPO on MPPs we described here could be one of the factors underlying the development of adverse side effects and co-morbidities during long-term EPO use in the clinics and associations of high EPO levels with leukemias^13, 67, 68^. To translate these results to the clinics and understand the side effect of EPO treatment, further work is required to determine if HSPCs and erythroid progenitors like CFU-E are responding to the same dose and duration of EPO exposure.

## Materials and Methods

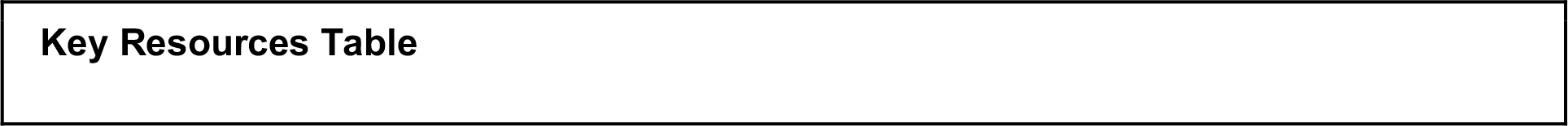
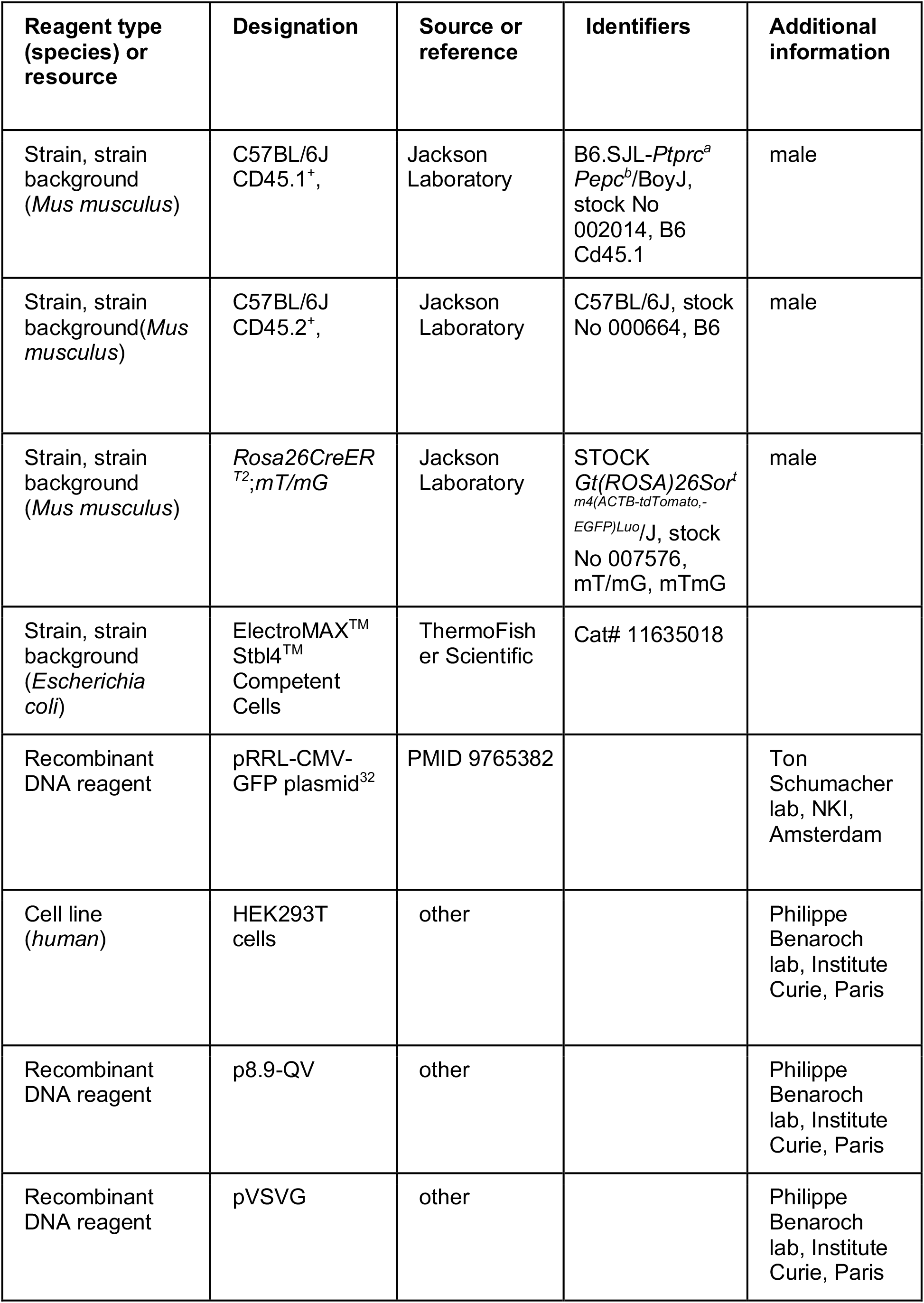
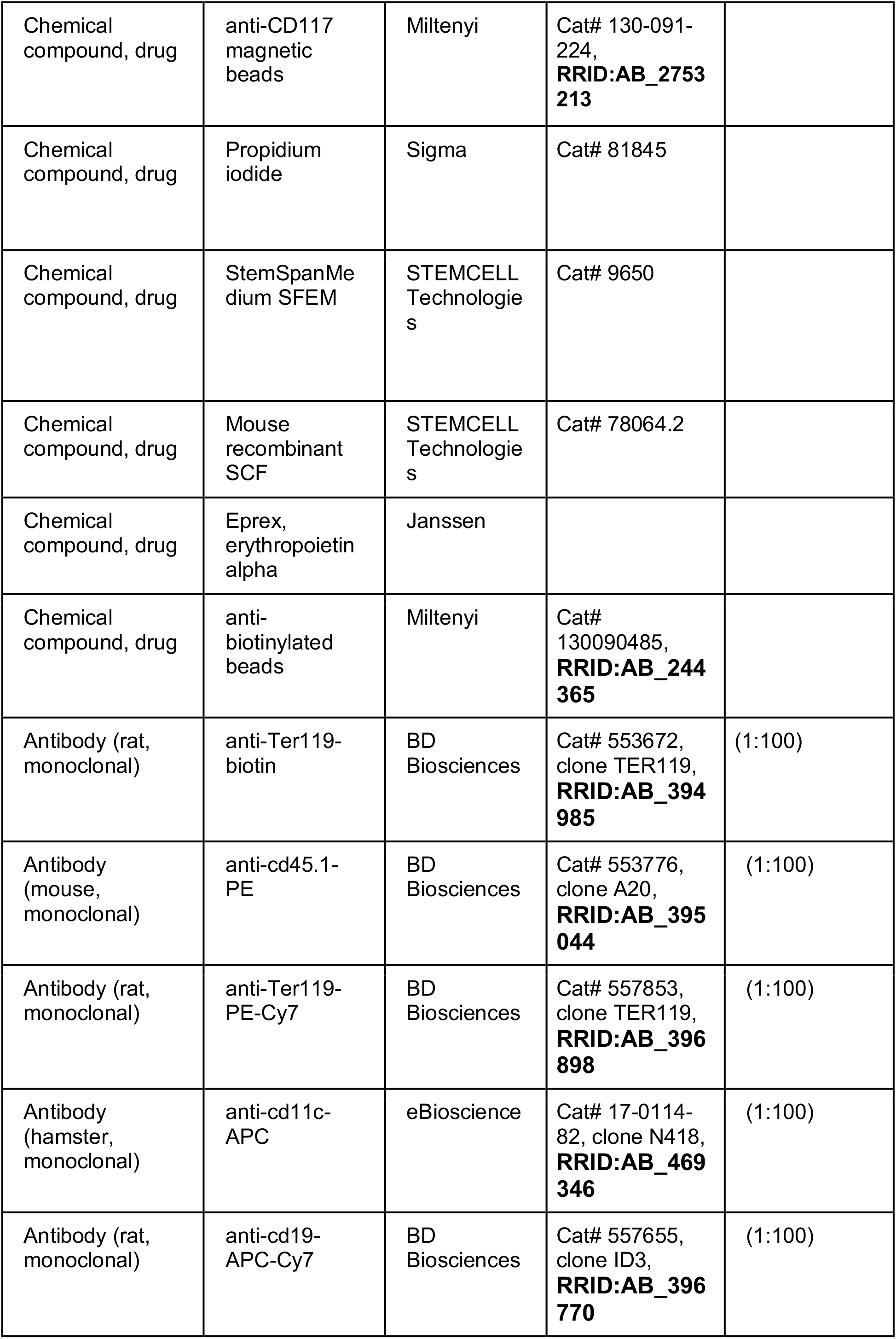
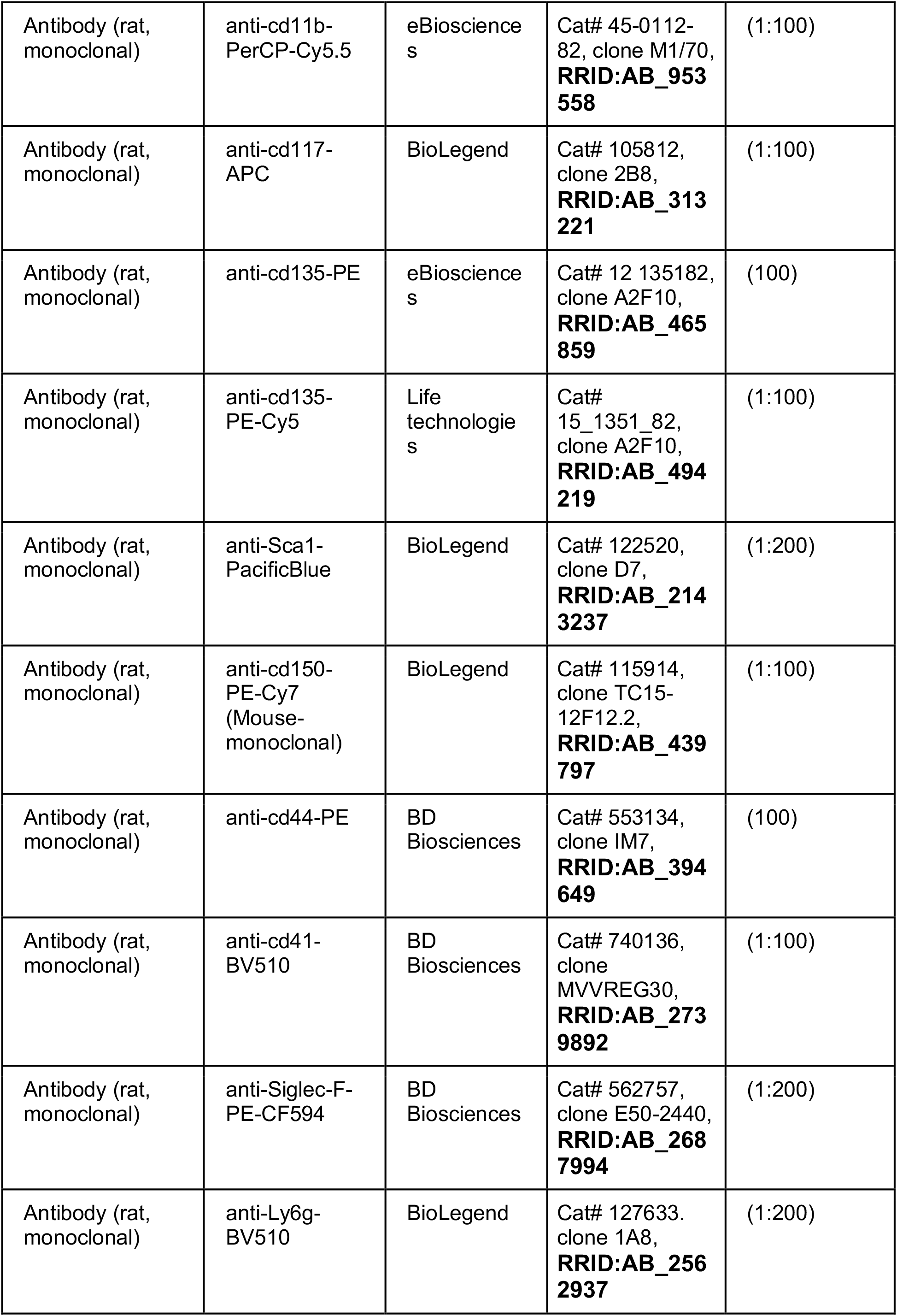
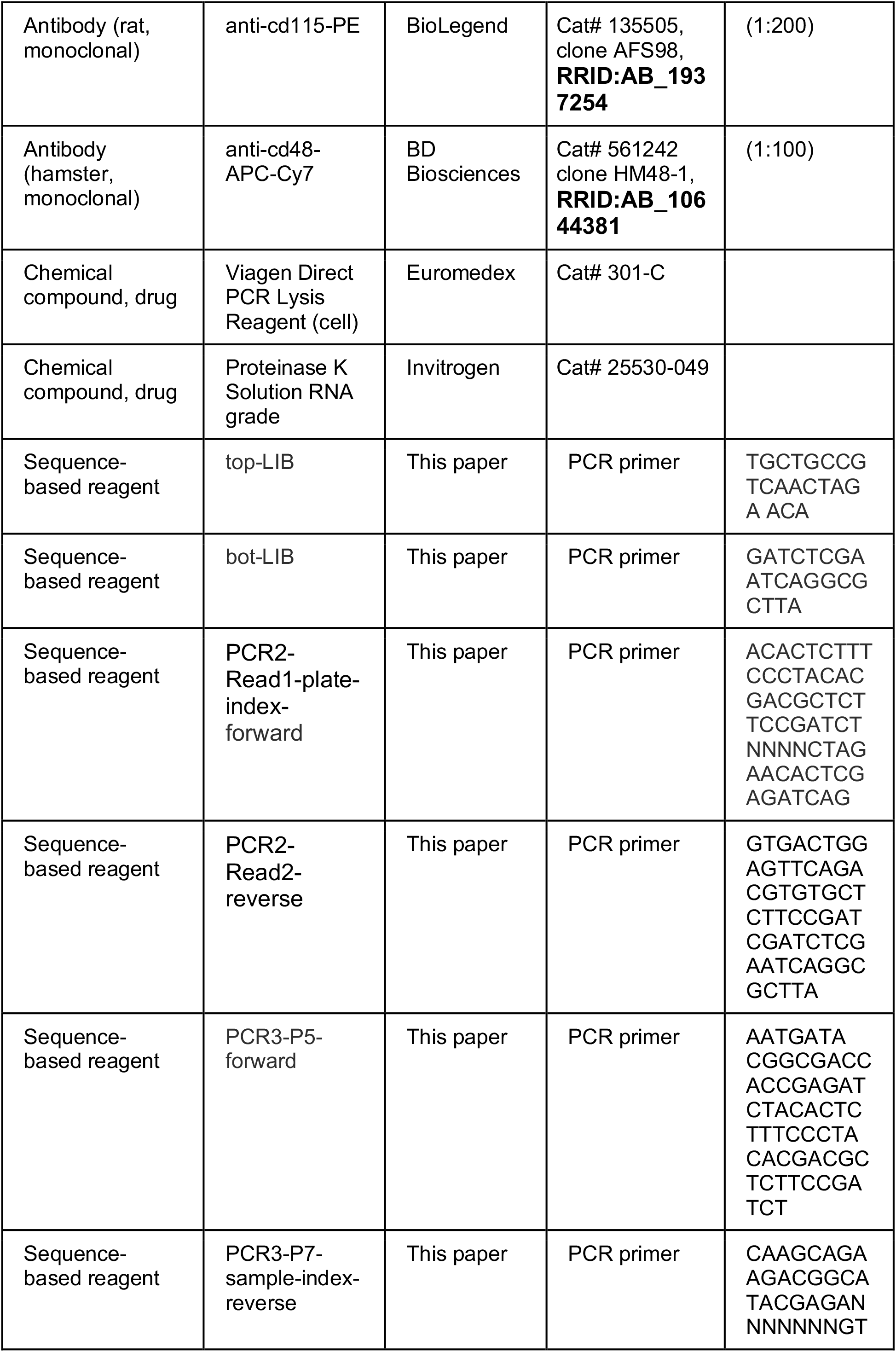
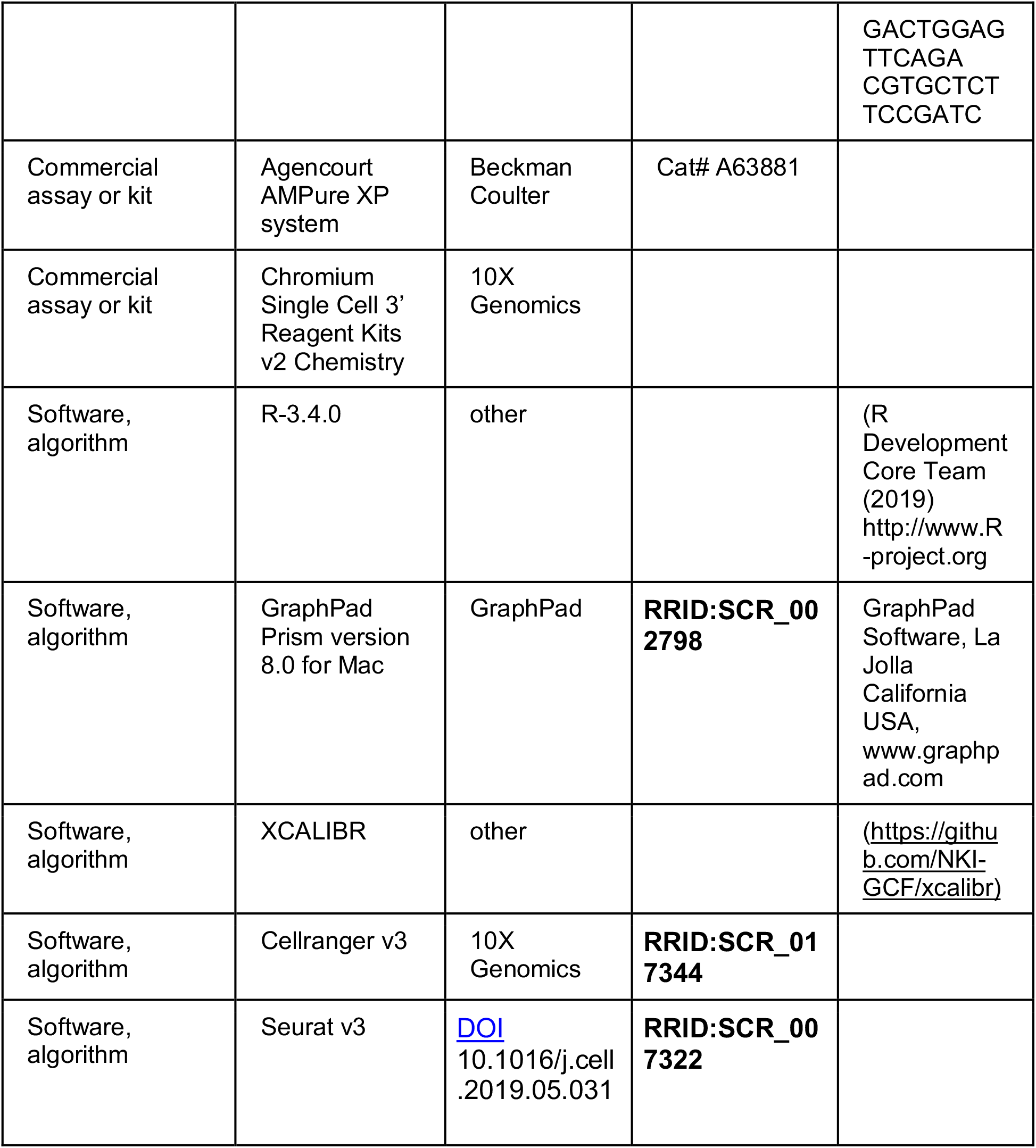

### Mice

Male C57BL/6J CD45.1^+^, C57BL/6J CD45.2^+^, and *Rosa26CreER^T^*^2^;*mT/mG* mice from Jackson Laboratory or bred at Institute Curie aged between 7-13 weeks were used in all experiments. All procedures were approved by the responsible national ethics committee (APAFIS#10955-201708171446318 v1).

### Barcode library, barcode reference list and lentivirus production

The LG2.2 barcode library is composed of a DNA stretch of 180 bp with a 20 bp “N”-stretch. DsDNA was generated by 10 PCR rounds and cloned into the XhoI-EcoRI site of the lentiviral pRRL-CMV-GFP plasmid^32^. Subsequently ElectroMaxStbl4 cells were transformed, and >10,000 colonies picked for amplification by Maxiprep. To create the barcode reference list (https://github.com/PerieTeam/Eisele-et-al<u>.-</u>), barcode plasmids were PCR amplified twice in duplicate and sequenced as described below. Sequencing results were filtered for barcode reference list generation as previously described in^33^. Lentiviruses were produced by transfecting the barcode plasmids and p8.9-QV and pVSVG into HEK293T cells in DMEM-Glutamax (Gibco) supplemented with 10% FCS (Eurobio), 1% MEM NEAA (Sigma), and 1% sodium pyruvate (Gibco) using Polyethyleneimine (Polysciences). Supernatant was 0,45 um filtered, concentrated by 1,5 h ultracentrifugation at 31,000g and frozen at –80°C. HEK293T were tested for mycoplasma contamination every 6 months.

### HSPC and MPP2 isolation, barcoding, EPO treatment, and transplantation

Isolation and labeling of cells with the barcoding library was performed as described in^33^. Briefly, after isofluorane aneasthesia and cervical dislocation, bone marrow cells were isolated from femur, tibia and iliac bones of mice by flushing, and C-Kit^+^ cells were enriched with anti-CD117 magnetic beads on the MACS column system (Miltenyi). Cells were stained for C-Kit, Flt3, CD150, Sca-1 (Key Resources Table) Propidium iodide (PI) (Sigma) (1:5,000), and if appropriate CD48. Lineage staining was not performed after C-Kit+ MACS enrichment for transplantation cohorts. For HSPC cohorts, HSPCs (Figure 1 – figure supplement 1a) were sorted and transduced with the barcode library in StemSpanMedium SFEM (STEMCELL Technologies) with 50 ng/ml mSCF (STEMCELL Technologies) through 1,5 h of centrifugation at 300 g and 4,5 h incubation at 37°C to obtain 10% barcoded cells. After transduction, cells were incubated with human recombinant EPO (Eprex, erythropoietin alpha, Janssen) at a final concentration of 1,000 or 160 ng/ml or PBS for 16 h at 37°C. After the incubation, the cells were transplanted by tail vein injection in recipient mice 6 Gy sub-lethally irradiated 3 hours before on a CIXD irradiator. Mice were allocated to groups of 4-5 mice for each condition randomly without masking. When indicated, cells were injected together with additional EPO (133 ug/kg). On average 2,600 cells (Mean 2,684 cells +/- 175 cells) were injected in the tail vein of each mouse. For the MPP2 cohort, MPP2 and CD48^-^ HSPCs (Figure 7 – figure supplement 1a) were sorted. Both populations were cultured alike, but only MPP2 were transduced with the barcode library and treated with 1,000 ng/ml recombinant EPO as described above. After the culture, barcoded MPP2 and un-barcoded CD48^-^ HSPCs were mixed at a ratio of 32/45 (to be as close as possible to the original ratio of both populations in the HSPCs) and transplanted as described above. A FACSAria^TM^ (BD Biosciences) was used for sorting. FACSDiva^TM^ software (BD Biosciences) was used for measurements and FlowJo^TM^ v.10 (TreeStar) for analysis.

### Cell progeny isolation for barcode analysis

Spleens were mashed and both blood and spleen cells were separated based on Ter119 using a biotinylated anti-Ter119 antibody (Key Resources Table) and anti-biotinylated beads on the MACS column system (Miltenyi). Ter119^+^ cells were stained for Ter119 and CD44^34^. Ter119^-^ cells were stained for CD45.1 CD11b, CD11c, CD19, and if appropriate CD115, Siglec-F, and Ly6G (Key Resources Table). Bone marrow cells were flushed from bones and enriched for C-Kit^+^ cells as above. When appropriate the C-Kit^-^ fraction was further separated based on Ter119 and stained as above. C-Kit^+^ cells were stained for C-Kit, Flt3, CD150, Sca-1, and if appropriate CD41 (Key Resources Table), and PI (1:5000) as described above. For analyzed and/or sorted populations see Figure 1 – figure supplement 1. Populations were only sorted for mice with an engraftment (donor cells percentage) of above 5% in spleen, bone and blood.

### Lysis, barcode amplification and sequencing

Sorted cells were lysed in 40 μl Viagen Direct PCR Lysis Reagent (cell) (Euromedex) supplemented with 0,5 mg/ml Proteinase K Solution RNA grade (Invitrogen) at 55°C for 120 min, 85°C for 30 min, 95°C for 5 min. Samples were then split into two replicates, and a three-step nested PCR was performed to amplify barcodes and prepare for sequencing. The first step amplifies barcodes (top-LIB (5′TGCTGCCGTCAACTAGA ACA-3′) and bot-LIB (5′GATCTCGAATCAGGCGCTTA-3′)). A second step adds unique 4 bp plate indices as well as Read 1 and 2 Illumina sequences (PCR2-Read1-plate-index-forward 5’ACACTCTTTCCCTACACGACGCTCTTCCGATCTNNNNCTAGAACACTCGAGAT CAG3′ and PCR2-Read2-reverse 5’GTGACTGGAGTTCAGACGTGTGCTCTTCCGAT CGATCTCGAATCAGGCGCTTA3′). In a third step, P5 and P7 flow cell attachment sequences and one of 96 sample indices of 7 bp are added (PCR3-P5-forward 5’AATGATA CGGCGACCACCGAGATCTACACTCTTTCCCTACACGACGCTCTTCCGATCT3′ and PCR7-P7-sample-index-reverse 5’CAAGCAGAAGACGGCATACGAGANNNNNNNGTGACTGGAGTTCAGA CGTGCTCTTCCGATC3′) (PCR program: hot start 5 min 95°C, 15 s at 95°C; 30 s at 57.2°C; 30 s at 72°C, 5 min 72°C, 30 (PCR1-2) or 15 cycles (PCR 3)). Both index sequences (sample and plate) were designed based on^35^ such that sequences differed by at least 2 bp (https://github.com/PerieTeam/Eisele-et-al<u>.-</u>). To avoid lack of diversity at the beginning of the reads, at least 4 different plate indices were used for each sequencing run. Primers were ordered desalted, as high-performance liquid chromatography (HPLC) purified. During lysis and each PCR, a mock control was added. The DNA amplification by the three PCRs was monitored by the run on a large 2% Agarose gel. PCR3 products for each sample and replicate were pooled, purified with the Agencourt AMPure XP system (Beckman Coulter), diluted to 5 nM. and sequenced on a HiSeq system (Illumina) (SR-65bp) at Institute Curie facility with 10% of PhiX spike-in.

### Barcode sequence analysis

Sequencing results were filtered, and barcodes were categorized in progenitor classes as in^33^ and further explained on github (https://github.com/PerieTeam/Eisele-et-al<u>.-</u>). In brief, sequencing results were analyzed using R-3.4.0 (R Development Core Team (2019) http://www.R-project.org.), Excel, and GraphPad Prism version 8.0 for Mac (GraphPad Software, La Jolla California USA, www.graphpad.com). Reads were first filtered for perfect match to the input index- and common-sequences using XCALIBR (https://github.com/NKI-GCF/xcalibr) and filtered against the barcode reference list. Samples were then filtered for containing at least 5000 reads and normalized to 10^5^ per sample. Samples with a Pearson correlation between duplicates below 0.9 were discarded and barcodes present in one of the two replicates were set to zero. Samples with under 10 barcodes were filtered out, unless indicated in the figure legend. The mean of the replicates was used for further processing. When the mean percentage of barcodes shared between different sequencing runs was higher than within the same sequencing run for mice of a same transduction batch, reads below the read quartile of the mean percentage of barcodes shared between mice of a same transduction batch but sequenced on different sequencing runs were set to zero in order to equalize the barcode sharing between mice transplanted from a same transduction batch in different sequencing runs to the barcode sharing between mice within each sequencing run. After filtering, read counts of each barcode in the different cell lineages were normalized enabling categorization into classes of biased output toward the analyzed lineages using a threshold of 10% of barcode reads (Other thresholds in Figure 1 – figure supplement 3c and Figure 7 – figure supplement 1c). Statistics on barcoding results were performed using a permutation test as in ^36^. Significance of flow cytometry results was assessed using Student’s T test. Some mice were excluded from the analysis due to death before readout, or due to a donor cell engraftment <5%, as well as the filtering out of mice for which one or more cell subset samples did not pass the barcode data filtering steps as detailed above.

### ScRNAseq and analysis

ScRNAseq was performed using the 10X Genomics platform on one pool of HSPCs isolated from 8 mice, barcoded and culture with or without EPO for 16h *in vitro* as described above. Sequencing libraries were prepared using the Chromium Single Cell 3’ v2 kit and sequenced on a HiSeq system (Illumina) at Institut Curie NGS facility. Data was analyzed using Cellranger v3 (10X Genomics), Seurat v3^37^ and customized scripts. Raw sequencing reads were processed using Cellranger. To obtain a reads/cell/gene count table, reads were mapped to the mouse GRCm38.84 reference genome. scRNAseq analysis was performed using Seurat^37^. During filtering, Gm, Rik, and Rp genes were discarded as non-informative genes. Cells with less than 1,000 genes per cell and with a high percentage of mitochondrial genes were removed from downstream analyses. Following our filtering procedures, the average UMI count per cell was 5157, with mitochondrial genes accounting for 5% of this. The average number of genes detected per cell was 2337. Cell cycle annotation using the cyclone method from the scran R package showed that 2938 cells were in G1 phase, 233 cells were in G2M phase, and 127 cells were in S phase. No batch effect was detected between the EPO and no-EPO group, therefore no batch correction was applied. Data normalization was performed using the default Seurat approach and differentially expressed genes were determined using a logistic regression in Seurat. Unsupervised clustering was performed on the significant variable genes using the ten first principle component analysis followed by the non-linear dimensionality reduction technique UMAP^38^ (Figure 5 – figure supplement 2). Unsupervised Louvain clustering of the data was performed across a range of resolution parameters and the resolution value that led to the most stable clustering profiles was chosen^39^ (Figure 5 – figure supplement 2). Annotation of the clusters was obtained by mapping published signatures using the *AddModuleScore* method of Seurat. The signatures are defined in the following publications: dHSC, and MPP1 signatures were obtained from^40^. The MolO LT-HSC signature was taken from^41^ and the MPP2 and 4 signature was taken from^31^. An excel file listing the genes in these signatures is available on github (https://github.com/PerieTeam/Eisele-et-al<u>.-</u>). To identify EPO-responder cells in the EPO group, differential expression analysis was performed between control and EPO groups (Lists of DEGs are available at https://github.com/PerieTeam/Eisele-et-al<u>.-</u>). Subsequently, genes that were differentially expressed (adjusted p-value< 0.05) between EPO and control groups were transformed into an EPO response signature which when overlaid onto the UMAP based visualization was enriched only in a subset of the EPO group cells. Briefly this signature was obtained by taking the background corrected mean expression values of both the up and downregulated genes per cell as implemented in the *AddModuleScore* method of Seurat. Within each cell these two signature scores were used to create a composite EPO response score by subtracting the downregulated response from the upregulated response signature. Cells in the upper 90^th^ percentile with regards to the expression of the EPO response signature were labelled EPO responders.

To perform supervised cell type annotation, a reference map was generated from a published single cell sequencing dataset of 44,802 C-Kit^+^ cells from^42^. Preprocessing was performed using a scanpy pipeline^43^. Data was then visualized using the non-linear non-dimensionality reduction technique UMAP^38^. Annotation of the reference map was obtained by overlaying published signatures as above using the *AddModuleScore* method of Seurat and also known markers as *Flt3*, *slamf1*, and *Gata1* (Figure 5 – figure supplement 1b). For the erythroid progenitors these markers are *Gata1, Klf1, Epor, Gypa, Hba-a2, Hba-a1* (figure 5 – figure supplement 1a). Cells were mapped onto the reference map using a k-nearest neighbors mapping approach. Briefly, for each cell in the query dataset, the nearest neighbors in PCA space of the reference dataset were determined using the nn2 function of the RANN package and the mean UMAP 1 and 2 coordinates of the 10 nearest neighbors were taken as the reference point for the new cell of interest. To benchmark our mapping approach, cells from an independent dataset of erythroid progenitors ^10^ were used without additional pre-processing (Figure 5 – figure supplement 1a).

## Data sharing statement

Raw data are available at zenodo DOI 10.5281/zenodo.5645045. All codes to filter and process raw data, as well as filtered data are available at https://github.com/PerieTeam/Eisele-et-al.-. Contact author is.

## Acknowledgements

We thank Dr. T. Schumacher for discussion and lentiviral library production and Dr. K Duffy for advices on permutation testing. We thank the Institute Curie flow cytometry, next generation sequencing, and animal facility. We thank Fahima Di Federico from the UMR168 BMBC facility for amplifying the barcode plasmid pool.

## Figure supplements legends

**Figure 1 – figure supplement 1:**
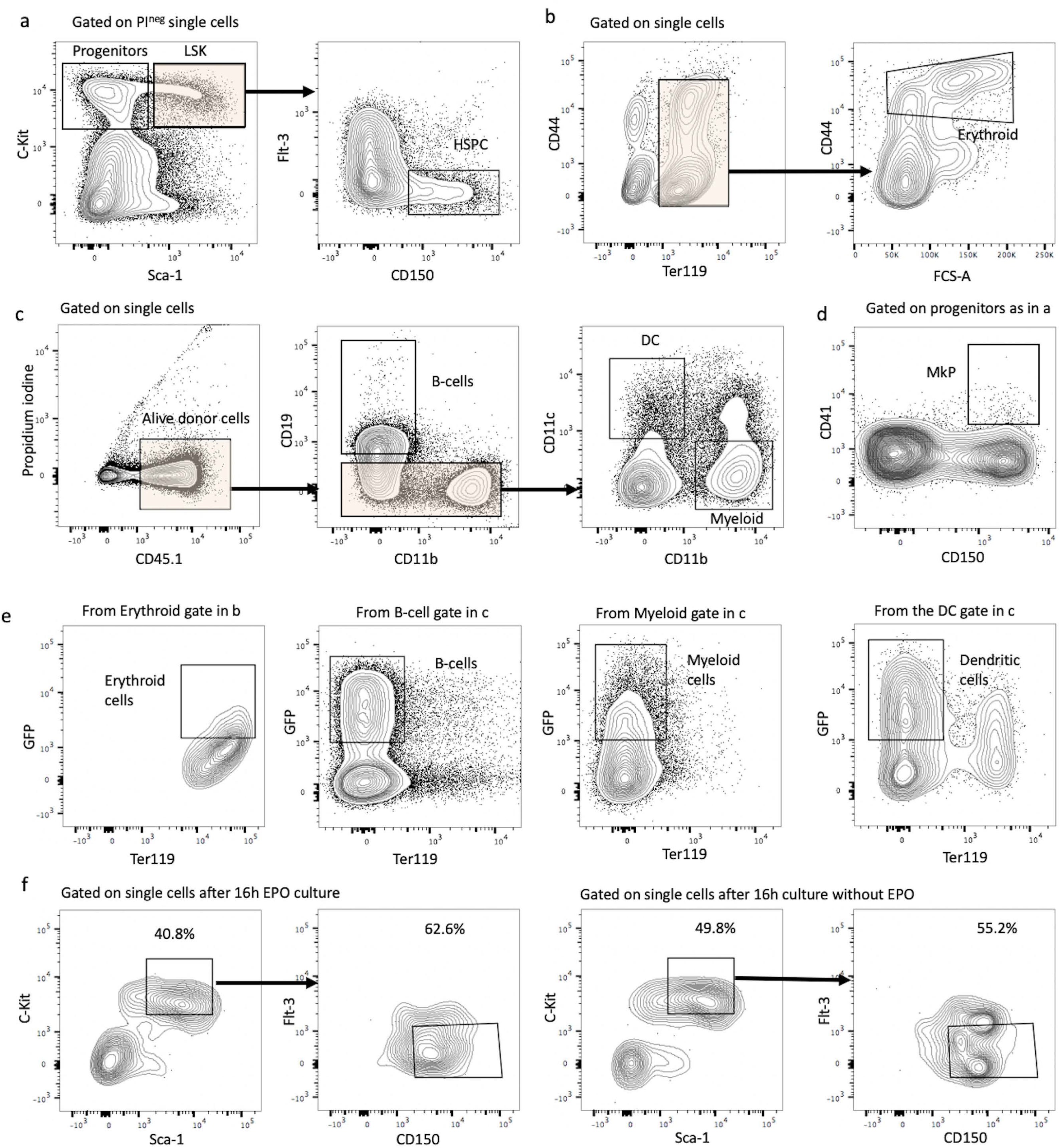
Gating strategies and HSPC marker expression after lentiviral transduction and *ex vivo* culture with or without EPO. **a**, HSPCs were gated as propidium iodide negative single C-Kit^+^ Sca-1^+^ Flt3^-^ CD150^+^ cells of C-Kit^+^ enriched bone marrow cells. **b**, Erythroblast cells were gated as Ter119^+^ CD44^+^ FSC^hi^ cells on Ter^+^ enriched cells. **c**, Gating strategy for B-cells (CD19^+^ CD11b^-^), dendritic cells (CD19^-^ CD11b^-^ CD11c^+^), and myeloid cells (CD119^-^ CD11c^-^ CD11b^+^) on Ter119^-^ live single donor cells. **d**, Gating for MkP (C-Kit^+^ Sca-1^-^ CD150^+^ CD41^+^) from C-Kit^+^ enriched bone marrow cells. **e**, Sort gating for GFP^+^ erythroid, myeloid, B-, and dendritic cells respectively used for barcoding analysis. **f**, Representative flow cytometry plots of sorted HSPC pool after 6 h lentiviral transduction and 16 h *ex vivo* incubation with or without EPO.

**Figure 1 – figure supplement 2:**
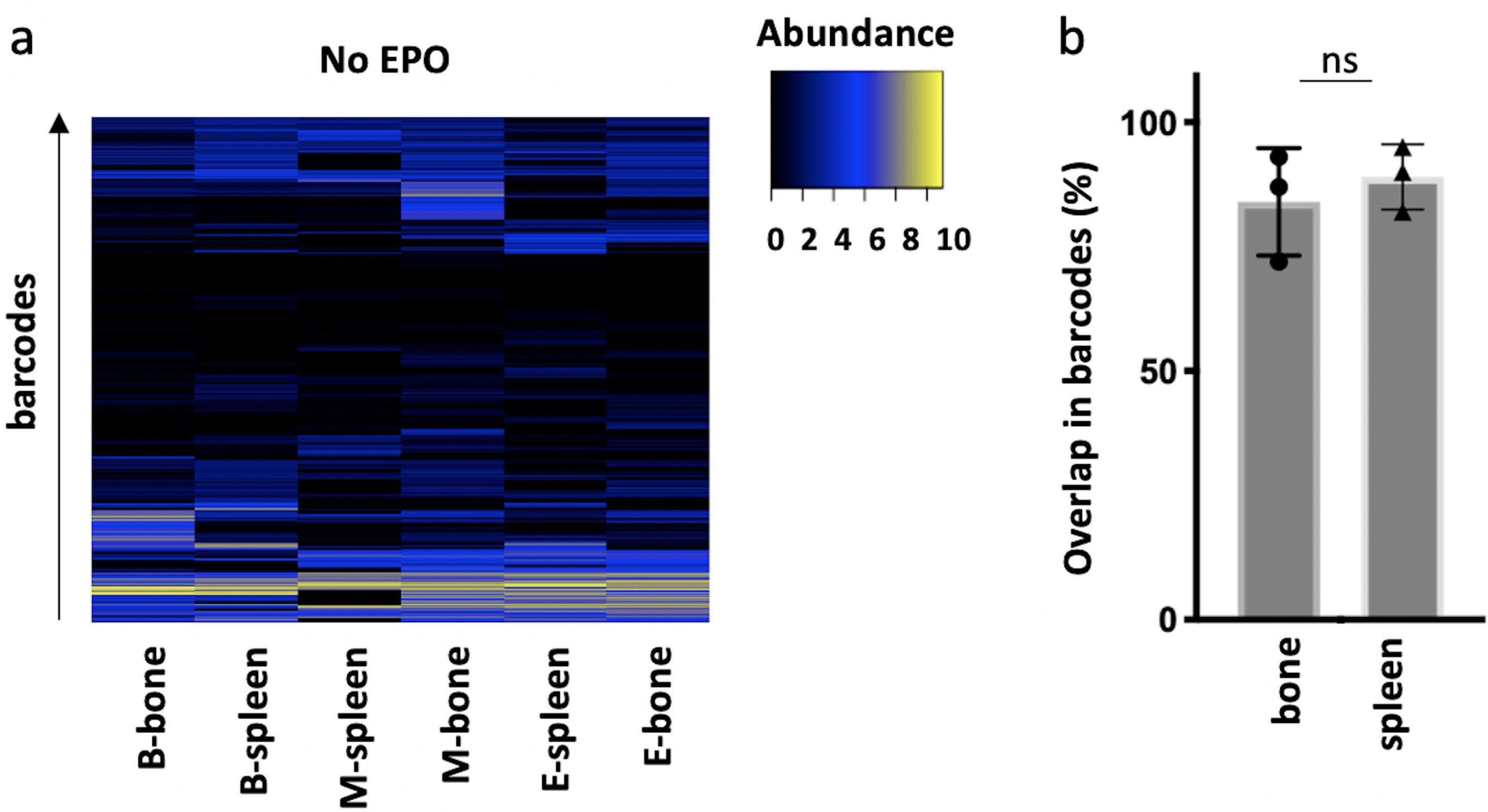
Correlations in barcoding profiles of spleen and bone. HSPCs were sorted from the bone marrow of donor mice, lentivirally barcoded, cultured *ex vivo* for 16 h and transplanted into sublethally irradiated mice. At week 4 post-transplantation, erythroid (E), B-cells (B), and the myeloid lineage (M) cells monocytes, eosinophils, neutrophils, and macrophages were sorted from the spleen and from bone and processed for barcode analysis. The myeloid lineage was merged according to the percentage of total donor myeloid each subset contributed as in Supplementary Figure 4a. **a**, Heatmaps showing the output of individual barcodes (rows) in different samples (columns) as indicated. Data is normalized by cell subset, log transformed, and clustered by complete linkage using Euclidean distance. No output is represented in black. **b**, The percentage of barcodes in spleen and bone detected in the respective other organ. The spearman rank correlation of barcodes in bone and spleen was for the B-, M- and E-lineage 0.81, 0,69 and 0,7 respectively. Shown are values from several animals (n= 3). For all bar graphs mean and S.D. between mice are depicted. Statistical significance tested using Mann-Withney U test p=0,05 for (b).

**Figure 1 – figure supplement 3:**
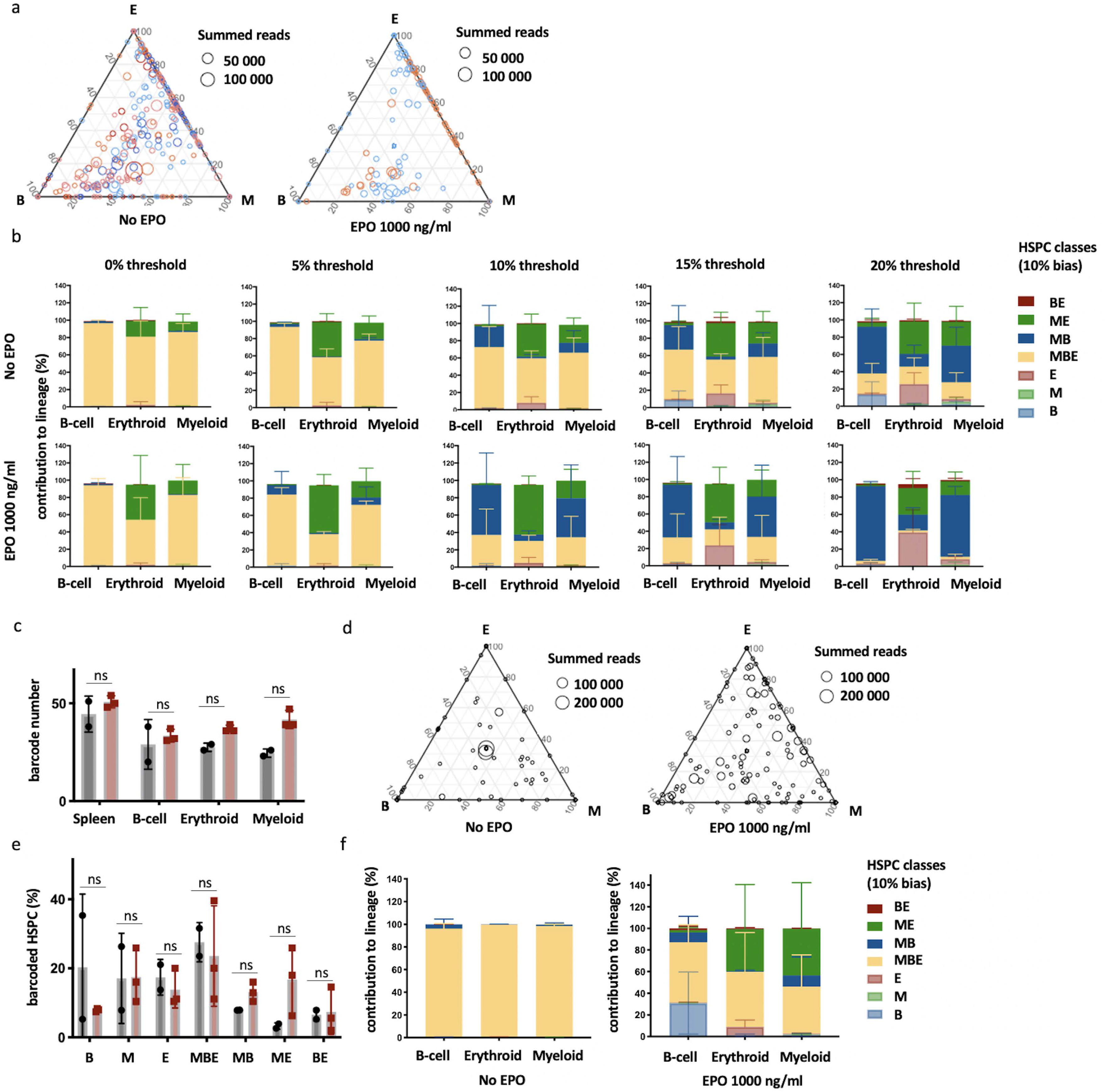
Characterization of lineage biases after transplantation of EPO-exposed HSPCs. **a**, Triangle plots from Figure 1e color coded by mice. **b**, Quantitative contribution of the classes to each lineage as in Figure 1 using different thresholds of 0%, 5%, 10%, 15% and 20%. **c-f**, Data for an additional experiment as in Figure 1. **c**, Number of barcodes retrieved in the indicated lineages at week 4 after transplantation in control and EPO group. **d**, Triangle plots showing the relative abundance of barcodes (circles) in the erythroid (E), myeloid (M), and B-cell (B) lineage with respect to the summed output over the three lineages (size of the circles) for control and EPO group. **e**, Proportion of HSPCs classified in the indicated lineage bias category, using a 10% classification threshold. **f**, Quantitative contribution of the classes as in f to each lineage. Shown are values from several animals (a-c, n=5 for control group and n=2 for EPO group (collected over one experiment), d-f, n= 2 for control, n= 3 for EPO group (collected over one experiment)). For all bar graphs mean and S.D. between mice are depicted. Statistical significance tested using Mann-Withney U test p=0,05 for (c, e).

**Figure 1 – figure supplement 4:**
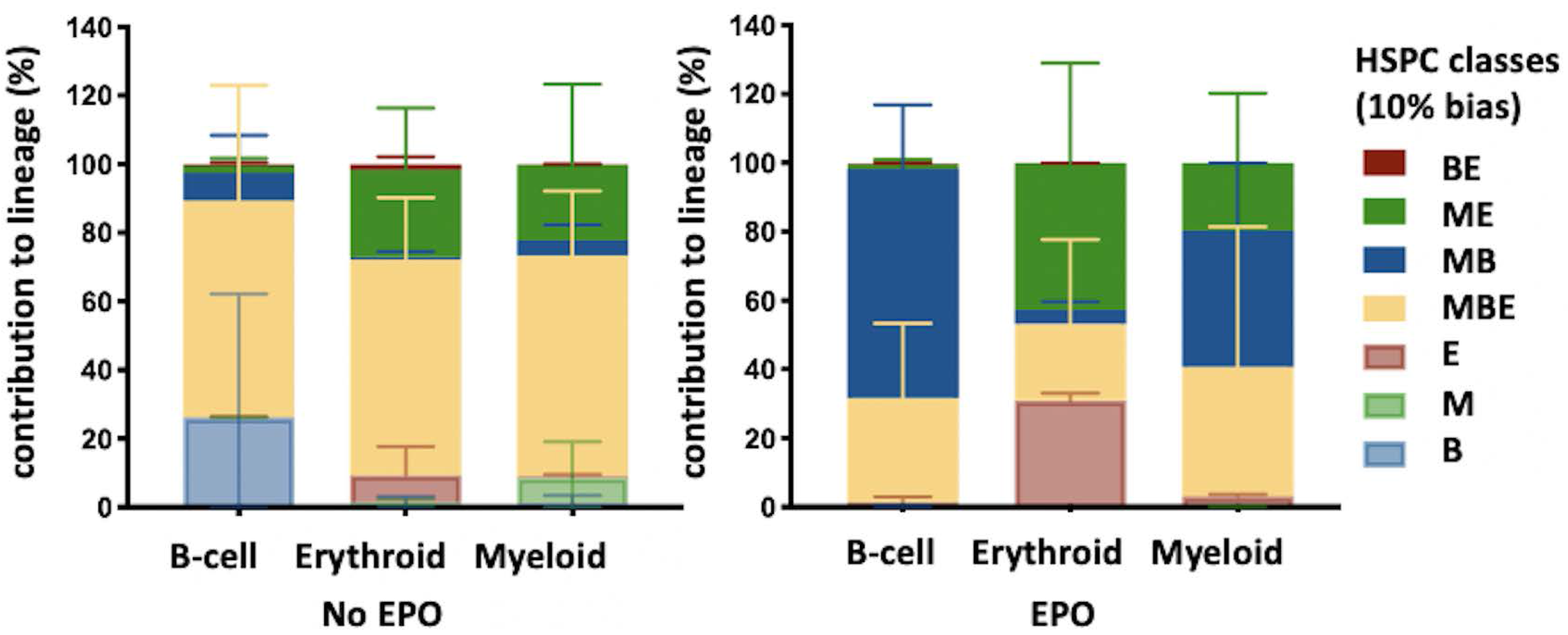
High output ME- and MB-biased HSPCs occur six weeks after transplantation of EPO-exposed HSPCs. HSPCs were sorted from the bone marrow of donor mice, lentivirally barcoded, cultured *ex vivo* with or without 1,000 ng/ml EPO for 16 h and transplanted into sublethally irradiated mice. At week 6 post-transplantation, the erythroid (E), myeloid (M), and B-cells (B) lineages were sorted from the spleen and processed for barcode analysis. Quantitative contribution of HSPCs classified by the indicated lineage bias, using a 10% threshold for categorization, to each lineage. Shown are values from several animals (n= 2 EPO, n= 4 control). Mean and S.D. between mice are depicted.

**Figure 2 – figure supplement 1:**
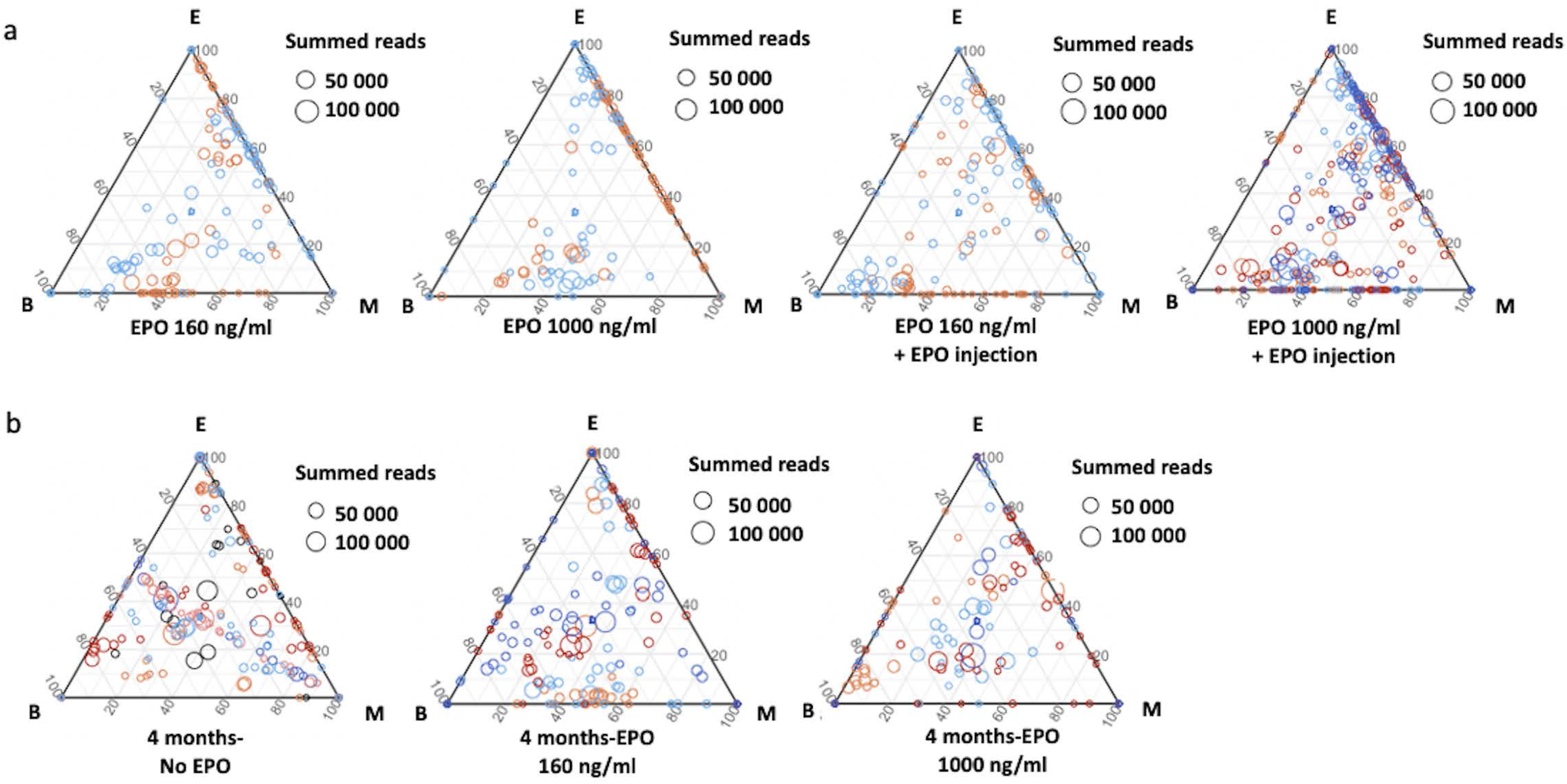
Variability in the effect of different EPO concentrations on clonality after HSPC transplantation at different timepoints. **a**, Triangle plots showing the relative abundance of barcodes (circles) in the erythroid (E), myeloid (M), and B-cell (B) lineage with respect to the summed output over the three lineages (size of circles) for control and EPO groups of Figure 2, color coded by mice. **b**, Same representation as in a for data of Figure 8.

**Figure 3 – figure supplement 1:**
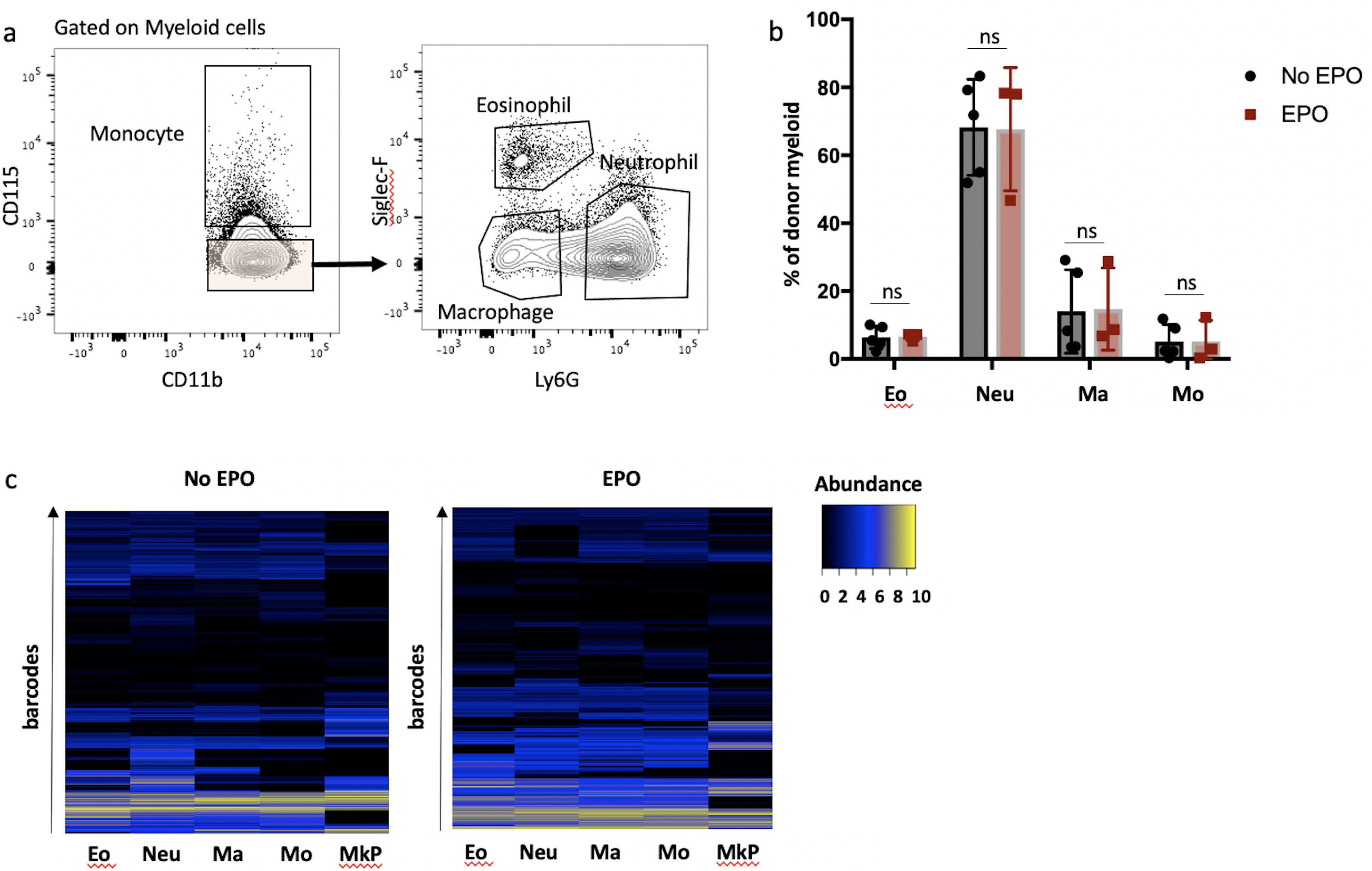
Production of macrophages, monocytes, neutrophils, eosinophils, and Megakaryocyte Progenitors (MkP) by HSPCs after EPO-exposure and transplantation. Same experimental protocol as in Figure 1 but the myeloid cells were subdivided into monocytes, eosinophils, macrophages and neutrophils, and MkP were sorted. **a**, Gating for detailed myeloid subsets on myeloid cells. Monocytes (Mo) were sorted as CD115^+^ cells, eosinophils (Eo) as CD115^-^ SiglecF^+^ Ly6G^+^, macrophages (Ma) as CD115^-^ SiglecF^-^ Ly6G^-^, and neutrophils (Neu) as CD115^-^ SiglecF^-^ Ly6G^+^ cells. **b**, The contribution of different cell types to the overall donor myeloid subset in control and EPO group. **c**, Heatmaps showing the output of individual barcodes (rows) in different samples (columns) as indicated. Data is normalized by cell subset, log transformed, and clustered by complete linkage using Euclidean distance. No output is represented in black. Shown are values from several animals (b, n=5 for control and n= 3 for EPO group c, n=3 for control and n=2 for EPO group (collected over two experiments)). For all bar graphs mean and S.D. between mice are depicted. Statistical significance tested using Mann-Withney U test p=0,05 for (b).

**Figure 5 – figure supplement 1:**
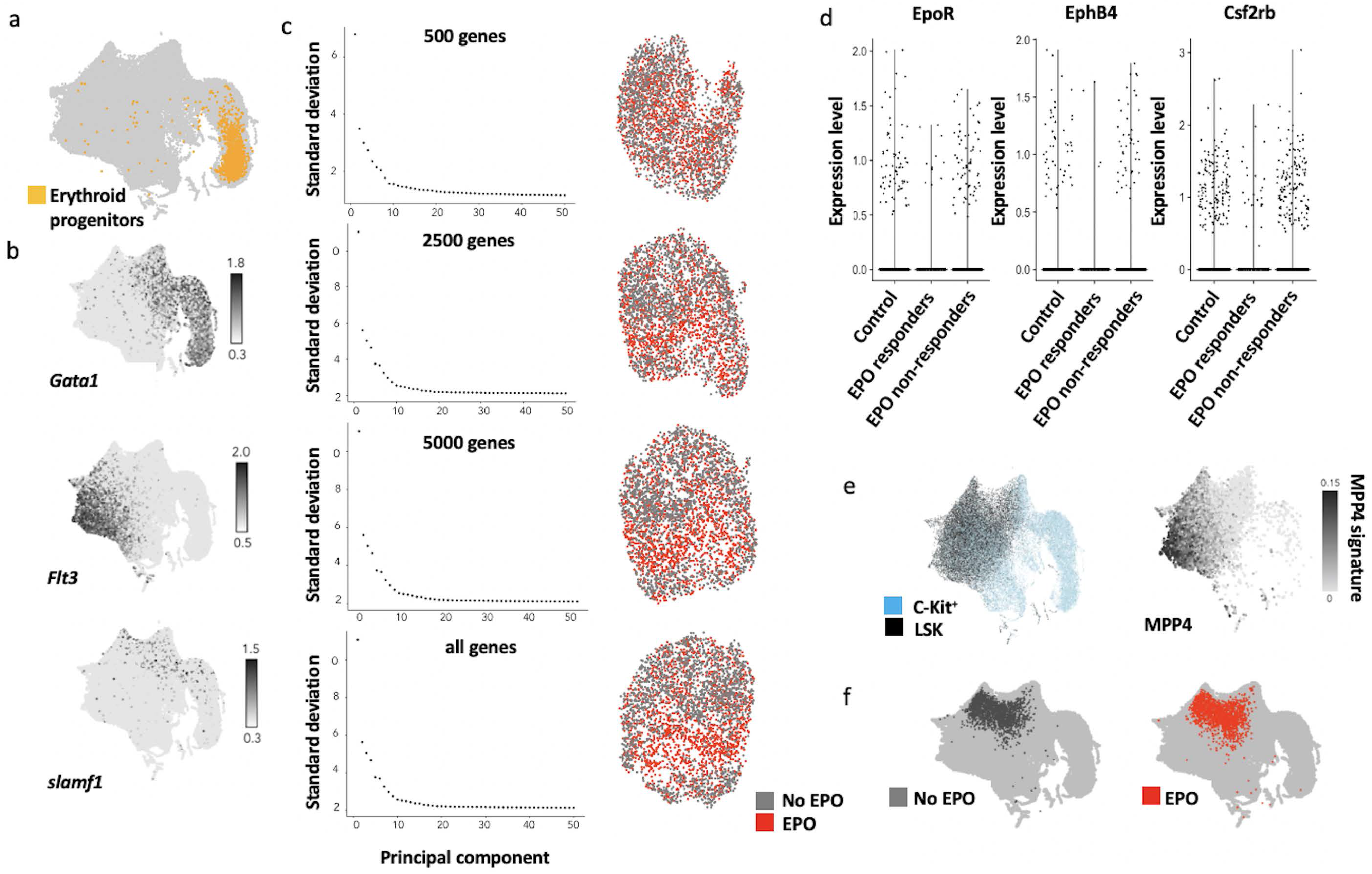
ScRNAseq analysis of control and EPO-exposed HSPCs. **a,** UMAP projection of the scRNAseq dataset from^42^ annotated with *flt3*, *CD150*, and *gata1* gene expression. **b**, Projecting of erythroid-biased progenitors from^10^ on UMAP projection of a. **c**, Robustness of the UMAP visualization and unsupervised clustering of the data in Figure 4c. The amount of variance explained by each principle component (left) and UMAP-based visualization using 10 PCA (right) for different number of genes. **d**, The expression of genes encoding known EPO receptors in each subgroup. **e,** Overview of the reference map using supervised cell type annotation of the dataset from^42^. On the right hand side we overlay the MPP4 signature defined by^31^ onto our reference map to facilitate celltype annotation **f**, HSPCs were sorted, barcoded and cultured *ex vivo* with or without 1,000 ng/ml EPO for 16 h, and analysed by scRNAseq using the 10X Genomics platform. Mapping of the transcriptomes of the 1,706 cells from control and 1,595 cells from EPO group obtained after quality control onto the reference map using a k-nearest neighbors mapping approach.

**Figure 5 – figure supplement 2:**
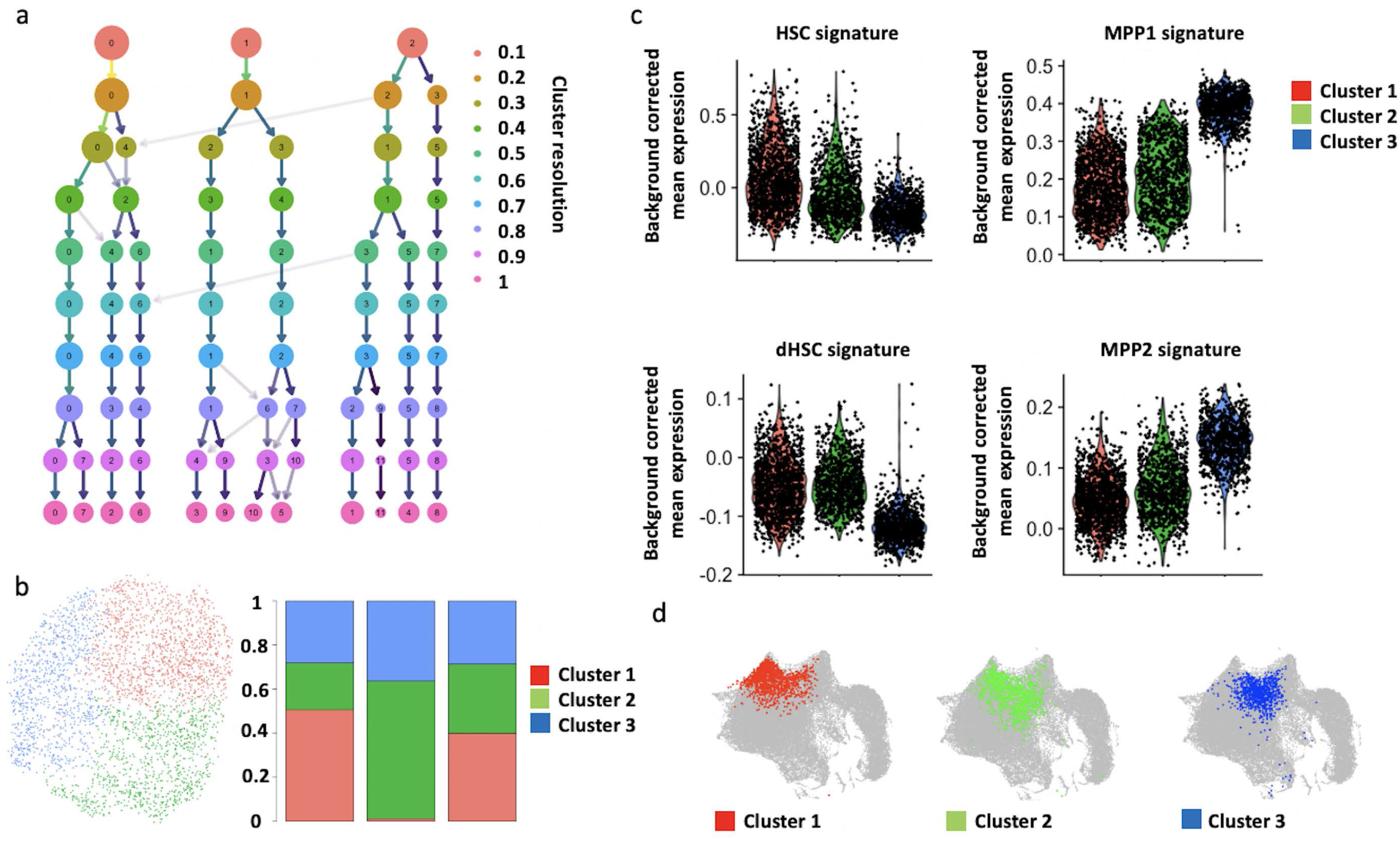
Unsupervised clustering of control and EPO-exposed HSPCs. **a,** Cluster stability analysis varying the resolution parameter of the Seurat clustering method. The significant variable genes using 10 PCA were used as input. **b**, UMAP visualization of the data in a using a clustering resolution of 0.1. with the proportion of each cluster as in the control, EPO-responder and non-responder subgroups. **c**, expression of published signatures of established celltypes used to annotate our clusters. All comparisons in signature expression between clusters were statistically significant (adjusted p-value < 0.05) as determined by a Kruskal-Wallis Test and Dunns Post-Hoc analysis. **d**, Nearest neighbor mapping of unsupervised clusters from b onto the reference map.

**Figure 7 – figure supplement 1:**
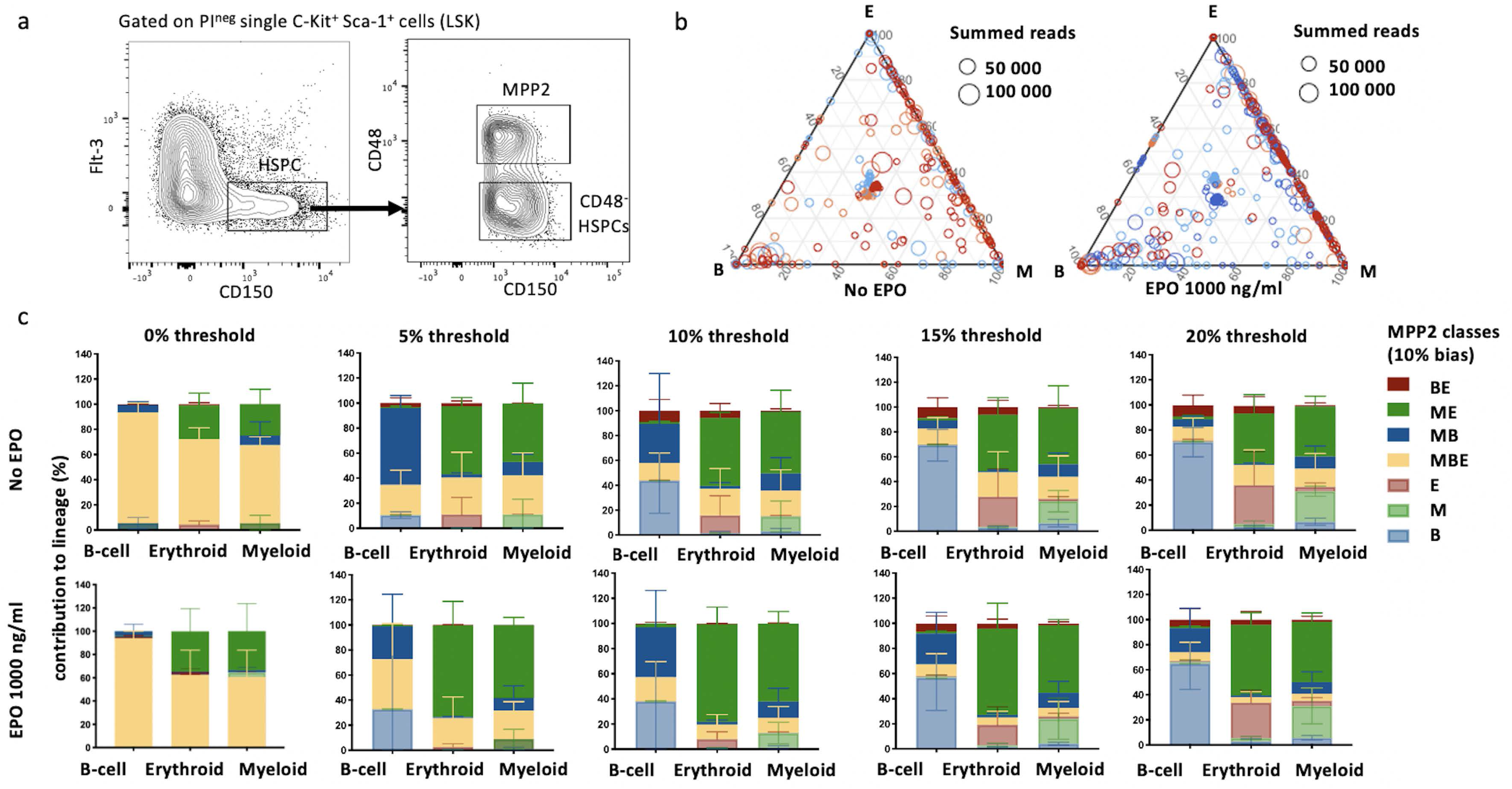
Characterization of lineage biases after transplantation of EPO-exposed MPP2. **a**, MPP2s and CD48^-^ HSPCs were gated as propidium iodide negative single C-Kit^+^ Sca-1^+^ Flt3^-^ CD150^+^ CD48^+^ (MPP2) and CD48^-^ (CD48^-^ HSPCs) cells of C-Kit^+^ enriched bone marrow cells. **b**, Triangle plots from Figure 7b color coded by mice. **c**, Quantitative contribution of the classes to each lineage as in Figure 7a using different thresholds of 0%, 5%, 10%, 15% and 20%. (a-c, n=5 for control group and n=2 for EPO group (collected over one experiment). Shown are values from several animals (n= 3 for control, n= 4 for EPO group (collected over one experiment)). For all bar graphs mean and S.D. between mice are depicted.

The article has one Supplementary File 1.

## Supplementary File legends

**Supplementary File 1: Permutation testing of changes in clonality after transplantation of EPO-exposed HSPCs.** Same data as in Figure 1 figure supplement 4 and Figure 1 figure supplement 3. HSPCs were cultured with EPO (1,000 ng/ml) for 16h. Barcodes in the erythroid (E), myeloid (M), B-lymphoid (B) lineage, dendritic cell (DC) and HSPCs, were analyzed four weeks after transplantation and categorized by bias using a 10% threshold. The output of MB and ME classified barcodes to the B, M, and E lineages was analyzed using a permutation test. By permutating the mice of control and EPO groups, the random distribution of this output was generated and compared to the real output difference between control and EPO group. A p-value was generated using permutation testing.

| Figure 1 figure supplement | Condition | p-value |  |  |  |
| --- | --- | --- | --- | --- | --- |
|  |  | MB in B | MB in M | ME in E | ME in M |
| 3 | HSPCs 1000 ng/ml repeat | 0.01 | 0.0083 | 0.0087 | 0.0087 |
| 4 | HSPCs 1000 ng/ml - 6 weeks timepoint | 0.01 | 0.01 | 0.0138 | 0.0185 |

